# Hmgb1 release kinetics shape its extracellular functions during regulated cell death

**DOI:** 10.64898/2026.05.15.725293

**Authors:** Shin Murai, Da Young Shin, Kenta Sumiyama, Kenta Moriwaki, Akira Muto, Kanako Takakura, Mai Yamagishi, Yoshitaka Shirasaki, Takumi Kanakogi, Sachiko Komazawa-Sakon, Tetuo Mikami, Masahiro Nishibori, Katsuhide Okunishi, Kenta Terai, Hiroyasu Nakano

## Abstract

Danger-associated molecular patterns (DAMPs), such as high mobility group protein B1 (HMGB1), are released from dying cells, yet the kinetics and functional consequences of their release remain unclear. Using Hmgb1–mCherry transgenic mice and live-cell imaging, we visualize Hmgb1 and interleukin-1β (IL-1β) secretion at single-cell resolution. Hmgb1 exhibits two distinct release kinetics: short-duration completed within 1 min and long duration spanning several minutes. Mathematical modeling demonstrates that short-duration release generates substantially higher local Hmgb1 concentrations and steeper gradients near the dying cell than long-duration release. Such rapid release may enable one or a few dying cells to produce sufficient Hmgb1 to stimulate neighboring cells, consistent with its role as an alarmin. In vivo imaging of cisplatin-induced kidney injury reveals an inverse correlation between intracellular Hmgb1 levels and monocyte infiltration. Together, these findings suggest that Hmgb1 release kinetics shape its extracellular functions during regulated cell death.

## Introduction

Dying cells release intracellular contents, which are referred to as damage-associated molecular patterns (DAMPs) ^1, 2^. DAMPs comprise a diverse range of molecules, including heat shock proteins (HSPs), high mobility group box 1 (HMGB1), ATP, and histones. Accumulating evidence indicates that these molecules exert pleiotropic functions, which are activated in a context-dependent manner ^1, 2^. Apoptotic cells maintain membrane integrity until the late stages, thereby limiting the release of large DAMPs. In contrast, other forms of regulated cell death—such as pyroptosis, necroptosis, and ferroptosis—facilitate the massive release of DAMPs through early membrane rupture ^3, 4^. Despite these insights, the precise spatiotemporal dynamics of DAMP release in vivo remain poorly understood.

HMGB1 is a nuclear protein that binds to chromatin ^5, 6^. HMGB1 is released from dying cells undergoing processes such as necroptosis, pyroptosis, and ferroptosis, likely through membrane rupture, and subsequently elicits inflammatory responses ^1, 2^. HMGB1 promotes the production of inflammatory cytokines by binding to TLR4 or RAGE, whereas it also exhibits chemokine-like activity through interactions with CXCL12 ^7, 8^.

We previously developed a F□rster resonance energy transfer (FRET) biosensor that can specifically monitor necroptosis termed the sensor for MLKL activation based on FRET (SMART) ^9^. Using this newly developed tool, we can visualize necroptosis both in vitro and in vivo ^9, 10^. To visualize DAMP release at single-cell resolution, we adapted live-cell imaging of secretion activity (LCI-S). This system utilizes an antibody-based sandwich ELISA combined with total internal reflection fluorescence (TIRF) microscopy ^11, 12^. By integrating LCI-S with SMART technology, we simultaneously monitored necroptosis execution and Hmgb1 release ^9^, revealing two distinct modes: a rapid, short-lived burst (within minutes) and a slower, sustained release (over 100 min). However, whether these modes have distinct biological functions remains unclear. Because the term “burst” in diffusion-based modeling denotes an instantaneous release at *t* = 0—an idealized condition unlikely to occur in biological systems ^13^—we hereafter refer to these patterns as short-duration and long-duration release to distinguish biological kinetics from mathematical terminology.

In the present study, we generate Tg mice expressing Hmgb1–mCherry to simultaneously visualize the release of Hmgb1–mCherry and IL-1β. Using these Tg mice and LCI-S technology, we analyze the release kinetics of Hmgb1 and IL-1β from macrophages during necroptosis and pyroptosis at single-cell resolution. Moreover, we apply a mathematical model to evaluate the biological outcomes of these two different release patterns. Finally, we analyze the expression of Hmgb1–mCherry in renal tubular epithelial (RTE) cells and the infiltration of immune cells by two-photon microscopy in a cisplatin-induced kidney injury model ^14, 15^. Collectively, these findings suggest that the intensity and duration of local signaling are governed by the specific mode of DAMP release. Furthermore, Hmgb1 release appears, at least in part, to drive the recruitment of circulating monocytes, linking cellular death kinetics to inflammatory cell infiltration.

## Results

### Generation of Hmgb1–mCherry Tg mice

To extend our previous analysis using a tumor cell line ^9^ to primary cells, such as macrophages, we generated Hmgb1**–**mCherry Tg mice. Hmgb1**–**mCherry was expressed across tissues at levels comparable to those of endogenous Hmgb1 (Fig. 1a). To confirm the expression of Hmgb1–mCherry in macrophages, we isolated peritoneal macrophages after thioglycolate injection and examined mCherry signals by flow cytometry. Nearly 90% of peritoneal macrophages from Tg mice were mCherry positive, whereas those from nontransgenic mice were mCherry negative (Fig. 1b and Supplementary Fig. 1). We next determined whether these macrophages were able to respond to canonical inflammasome or necroptotic stimuli. Cells treated with LPS plus nigericin (pyroptosis inducers) produced mature interleukin-1β (IL-1β) and the N-terminal fragment of Gsdmd (Gsdmd-N), whereas cells treated with BV6 plus zVAD (necroptosis inducers) did not (Fig. 1c).

**Figure 1.**
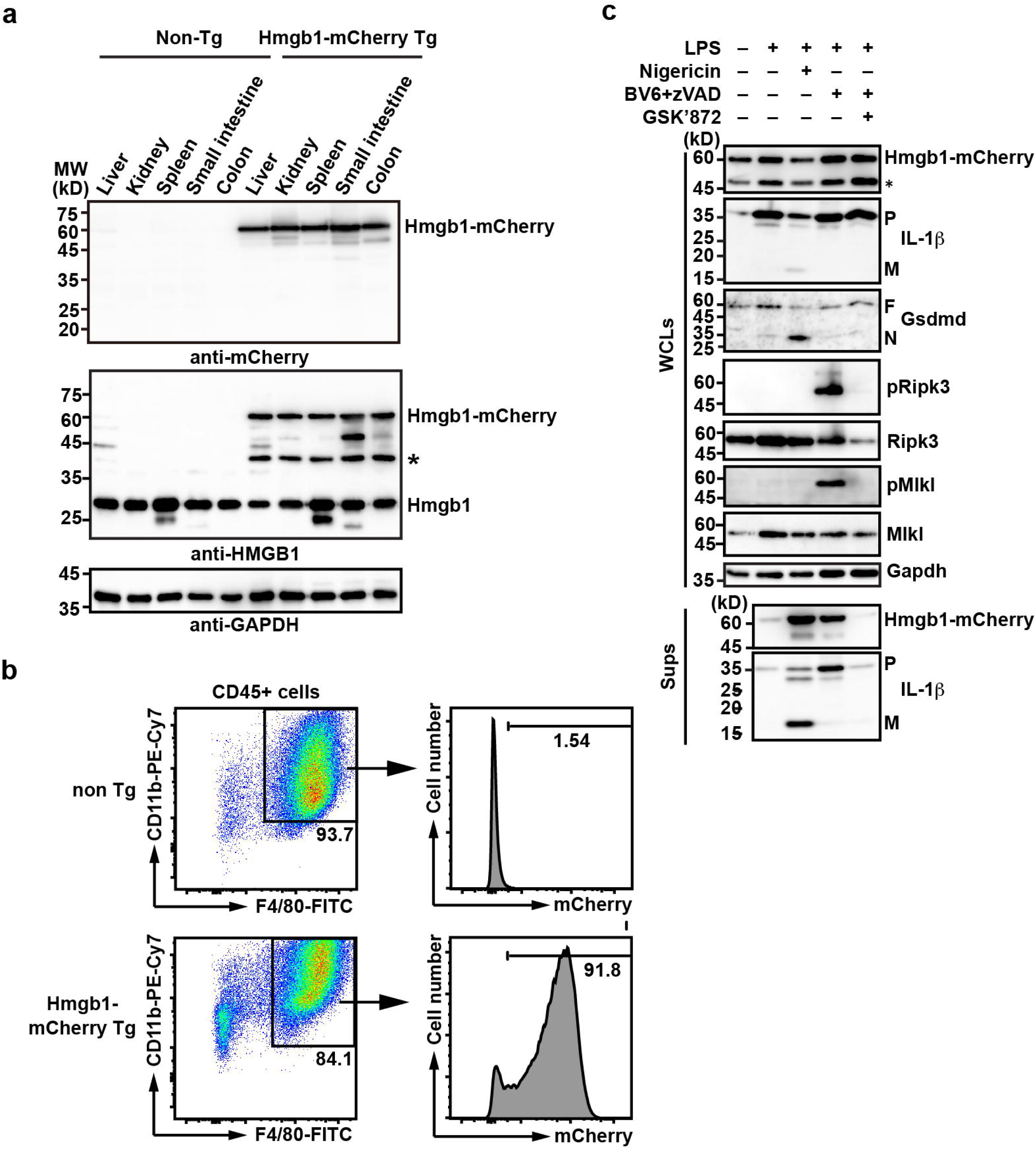
Generation of Hmgb1–mCherry Tg mice. **a** Hmgb1–mCherry was expressed in various murine tissues. Tissue extracts were prepared from the indicated organs of 8-week-old wild-type or Hmgb1–mCherry Tg mice. The expression of Hmgb1–mCherry was analyzed by Western blotting with the indicated antibodies. The asterisk indicates the cross-reactive band. **b** Expression of Hmgb1–mCherry in peritoneal macrophages. Mice of the indicated genotypes were intraperitoneally injected with thioglycollate, and peritoneal cells were subsequently recovered by washing the peritoneal cavity with ice-cold PBS on Day 4 after injection. Isolated cells were stained with the indicated antibodies and analyzed by flow cytometry. The percentages of CD11b^+^ F4/80^+^ cells (presumably peritoneal macrophages) are shown on the left. The percentages of mCherry-positive cells and their fluorescence intensities are shown as histograms on the right. **c** Peritoneal macrophages from Hmgb1–mCherry Tg mice were left untreated or primed with LPS (10 ng mL^−1^) for 3 h and then stimulated with nigericin (5 µM) for 1 h to induce pyroptosis or BV6 (1 μM) + zVAD (20 μM) to induce necroptosis in the absence or presence of the RIPK3 inhibitor GSK’872 (5 μM) for 4 h. The culture supernatants (Sups) or whole-cell lysates (WCLs) were analyzed by Western blotting with the indicated antibodies. P and M indicate the proform and mature form of IL-1β, respectively; F and N indicate the full-length and N-terminal fragment of Gsdmd, respectively. The asterisks indicate cross-reactive bands. All the results are representative of two or three independent experiments. Source data are provided as Supplementary Data 1.

LPS/nigericin induced the release of full-length Hmgb1**–**mCherry together with mature IL-1β into the culture supernatant, while BV6/zVAD induced the release of Hmgb1**–**mCherry and pro-IL-1β (Fig. 1c). These results demonstrate that peritoneal macrophages from Hmgb1**–**mCherry Tg mice faithfully recapitulate the responses of wild-type macrophages and provide a robust system for analyzing Hmgb1**–**mCherry and IL-1β release using LCI-S.

### Distinct release modes of Hmgb1 and IL-1β during pyroptosis

Previous studies have established that in pyroptotic macrophages, IL-1β is released via GSDMD-mediated pore formation, whereas Hmgb1 release is driven by NINJ1-induced membrane rupture ^16, 17, 18^. To visualize the release of Hmgb1**–**mCherry and endogenous IL-1β simultaneously from macrophages, we stimulated peritoneal macrophages with LPS plus nigericin. Interestingly, ∼50% of the macrophages consistently released Hmgb1**–**mCherry alone, whereas ∼30% of the macrophages released both Hmgb1**–**mCherry and IL-1β (Fig. 2a). The remaining fraction released only IL-1β, which is somewhat consistent with the ∼10% of macrophages that did not express Hmgb1**–**mCherry (Fig. 1b). These results suggest that the extracellular release of each DAMP does not always occur simultaneously, depending on the cellular context. In addition, the fluorescence intensity of extracellularly released Hmgb1 was significantly higher in cells exhibiting co-release compared to those releasing Hmgb1 alone (Fig. 2b).

**Figure 2.**
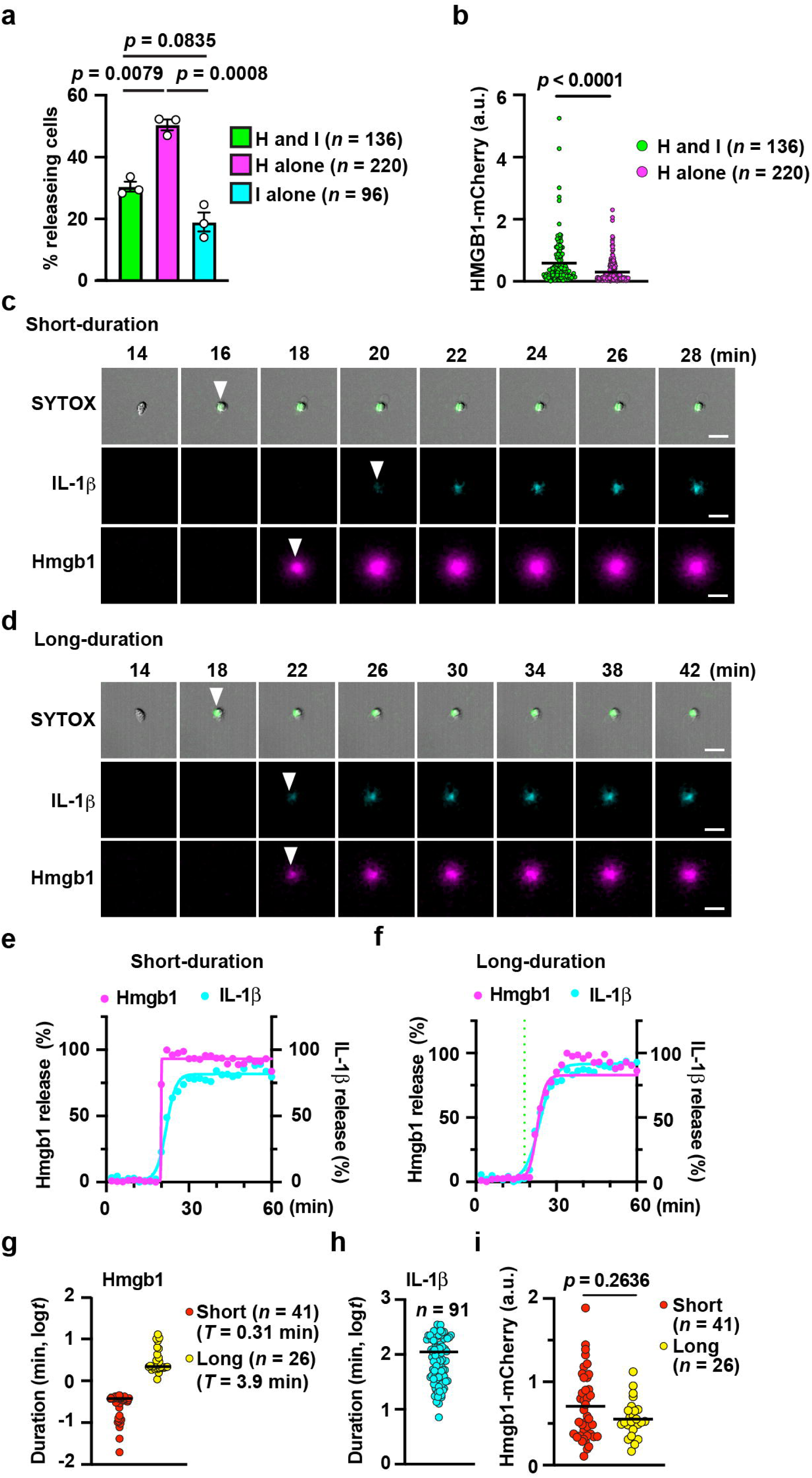
Distinct release modes of Hmgb1 and IL-1β during pyroptosis. Peritoneal macrophages isolated from Hmgb1–mCherry Tg mice were primed with LPS (10 ng mL^−1^) for 3 h and subsequently stimulated with nigericin (20 μM). The release of Hmgb1–mCherry and IL-1β was analyzed by LCI-S over 8 h. **a** Quantification of cells releasing both Hmgb1–mCherry and IL-1β (H and I), Hmgb1–mCherry alone (H alone), or IL-1β alone (I alone). Cells exhibiting each release pattern were counted, and the percentages were calculated relative to the total number of released cells. Results are mean ± SD from three independent experiments (H and I*, n* = 136 cells; H alone, 220 cells; I alone, 96 cells). **b** Relative Hmgb1–mCherry fluorescence intensities in H and I cells versus H alone cells. The fluorescence intensity of each individual cell was quantified by LCI-S. Each dot represents a single cell pooled from three independent experiments (H and I, *n* = 136 cells; H alone, *n* = 220 cells). Results are mean ± SD. a.u., arbitrary unit. **c–f** Representative image montages (**c, d**) and release kinetics (**e, f**) of short-duration (**c, e**) and long-duration (**d, f**) cells. The release of IL-1β (blue) and Hmgb1–mCherry (magenta), together with SYTOX uptake (green), was visualized by LCI-S. White arrows indicate the timing of SYTOX uptake and the onset of Hmgb1–mCherry and IL-1β release. Representative release kinetics of Hmgb1–mCherry (magenta) and IL-1β (blue) from short-duration (**e**) and long-duration cells (**f**) are shown. Time 0 indicates the addition of nigericin. **g** Dual modes of Hmgb1–mCherry release. The duration of Hmgb1–mCherry release was calculated as described in the Methods section and analyzed by *k*-means clustering, and two groups were identified: short-duration cells (*n* = 41) and long-duration cells (*n* = 26). **h** Single release mode of IL-1β. The release duration was calculated as in (**g**) and classified into a single cluster (*n* = 91 cells). The results in **g** and **h** are representative of three independent experiments. **i**, Relative Hmgb1–mCherry fluorescence intensities in short-duration versus long-duration cells. Data are presented as mean ± SD (short-duration, *n* = 41; long-duration, *n* = 26). a.u., arbitrary unit. Statistical significance was assessed by the one-way ANOVA followed by Tukey’s multiple comparison test (**a**) or Mann–Whitney *U* test (**b, i**). Representative results from three independent experiments are shown (**c–h**). Source data are provided as Supplementary Data 1.

We next examined the kinetics of Hmgb1 release. The duration of release was defined as the time interval between 5% and 95% of the maximal signal, determined by sigmoidal curve fitting. Membrane rupture was defined as the time at which SYTOX fluorescence intensity reached its peak, serving as a temporal marker for the completion of plasma membrane permeabilization. Consistent with our previous observations in L929 cells ^9^, Hmgb1–mCherry release from macrophages undergoing pyroptosis exhibited two distinct kinetic patterns: a short-duration mode terminating within ∼1 min and a long-duration mode persisting for several minutes (Fig. 2c–f, Supplementary Movies 1 and 2). To quantitatively classify these patterns, we performed *k*-means clustering on the log-transformed duration values (Supplementary Fig. 2) ^19^, with the number of clusters determined on the basis of statistical separation and biological interpretability. This analysis revealed two distinct clusters with mean durations of 0.31 and 3.89 min (Fig. 2g). In contrast, IL-1β release did not show clear separate patterns and was therefore treated as continuous (Fig. 2h). Additionally, the total amount of Hmgb1–mCherry released was comparable between short-duration and long-duration release modes (Fig. 2i).

Notably, direct comparison of Hmgb1 and IL-1β kinetics was not feasible because of their distinct detection formats: Hmgb1–mCherry is visualized immediately upon direct capture ^9^, whereas IL-1β was untagged and requires a sandwich-based approach involving a fluorescently labeled secondary antibody ^11^.

### Distinct release modes of Hmgb1 and IL-1β during necroptosis

Given that IL-1β is cleaved by caspase-1 and subsequently released through GSDMD pores, direct comparison of the release kinetics between IL-1β and HMGB1 is inherently complex, as HMGB1 is exclusively released following NINJ1-mediated plasma membrane rupture in pyroptotic cells ^18^. In contrast, in necroptotic cells, both IL-1β and HMGB1 are released upon MLKL-mediated membrane rupture. To address this difference, we examined the release kinetics of these proteins in cells undergoing necroptosis induced by BV6/zVAD following priming with LPS. Consistent with our observations in pyroptotic cells, approximately 40% of the cells released both Hmgb1–mCherry and IL-1β and approximately 40% released Hmgb1–mCherry alone, whereas ∼20% of cells released only IL-1β (Fig. 3a). The fluorescence intensity of the extracellularly released Hmgb1 from coreleasing cells was greater than of the cells that released only Hmgb1 (Fig. 3b). Under these conditions, Hmgb1–mCherry release from necroptotic cells exhibited a bimodal pattern, comprising short- and long-duration events (mean durations of 0.30 and 10.24 min, respectively), whereas IL-1β release followed a unimodal pattern (Fig. 3c–h, Supplementary Movies 3 and 4). The total amount of Hmgb1–mCherry released from long-duration cells was slightly lower than that from short-duration cells, although the difference was not substantial (Fig. 3i). These results indicate that Hmgb1–mCherry is released from macrophages undergoing pyroptosis and necroptosis via two different modes.

**Figure 3.**
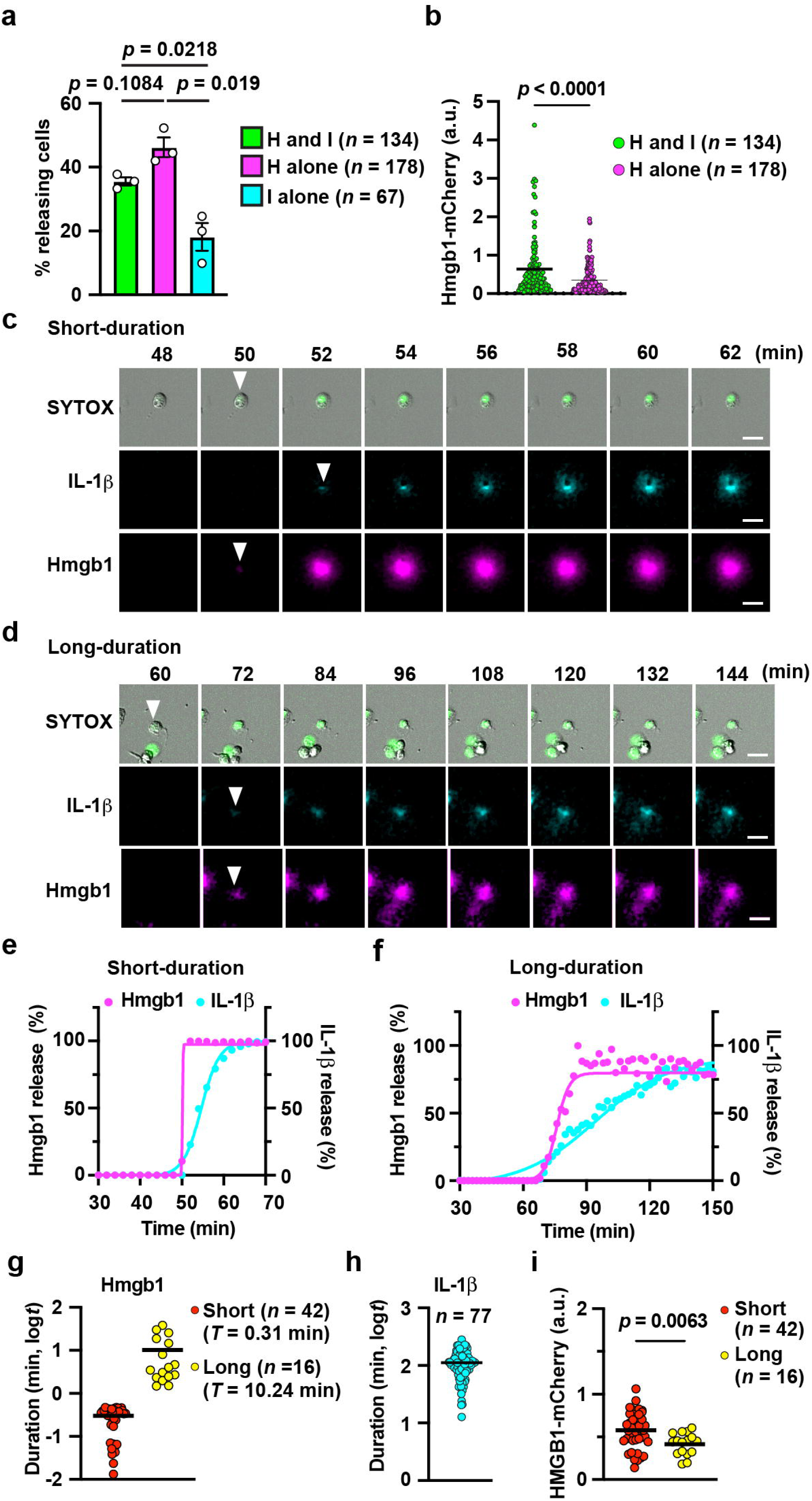
Distinct release modes of Hmgb1 and IL-1β during necroptosis. Peritoneal macrophages isolated from Hmgb1–mCherry Tg mice were primed with LPS (10 ng mL^−1^) for 3 h and subsequently stimulated with BV6 (1 μM) + zVAD (20 μM). The release of Hmgb1–mCherry and IL-1β was monitored by LCI-S over 8 h. **a** Quantification of cells releasing both Hmgb1–mCherry and IL-1β (H and I), Hmgb1–mCherry alone (H alone), or IL-1β alone (I alone), based on the release patterns as shown in Fig. 2a. Results are mean ± SD from three independent experiments (H and I, *n* = 134 cells; H alone*, n* = 178 cells; I alone*, n* = 67 cells). **b** Relative Hmgb1–mCherry fluorescence intensities quantified by LCI-S. Each dot represents a single cell pooled from three independent experiments (H and I, *n* = 134 cells; H alone*, n* = 178 cells). Data are shown as mean ± SD. a.u., arbitrary unit. **c–f** Representative image montages (**c, d**) and release kinetics (**e, f**) of short-duration and long-duration cells. The release of Hmgb1–mCherry (magenta) and IL-1β (blue), together with SYTOX uptake (green), was visualized by LCI-S. White arrows indicate the timing of SYTOX positivity and the onset of IL-1β and Hmgb1–mCherry release. Representative release kinetics of Hmgb1–mCherry (magenta) and IL-1β (blue) from short-duration (**e**) and long-duration cells (**f**) are shown. Time 0 indicates the addition of nigericin. **g** Dual modes of Hmgb1–mCherry release. The duration of Hmgb1–mCherry release was calculated as described in the Methods section and analyzed by *k*-means clustering, and two groups were identified: short-duration cells (*n* = 42) and long-duration cells (*n* = 16). **h** Single release mode of IL-1β. The release duration was calculated as in (**g**) and classified into a single cluster (*n* = 77 cells). Representative data from three independent experiments are shown. **i** Relative Hmgb1–mCherry fluorescence intensities in short-duration versus long-duration cells. Data are presented as mean ± SD (short-duration, *n* = 42; long-duration, *n* = 16). a.u., arbitrary unit. Statistical significance was assessed by the one-way ANOVA with Tukey’s multiple comparison test (**a**), Mann–Whitney *U* test (**b**), or two-tailed unpaired Student’s *t*-test (**i**). Representative results from three independent experiments are shown (**c-h**). Source data are provided as Supplementary Data 1.

### Distinct release sites of Hmgb1 and IL-1β during pyroptosis

Because mature IL-1β and larger DAMPs such as HMGB1 are released through GSDMD pores and NINJ1-mediated membrane rupture, respectively ^18^, it remains unclear whether these molecules originate from the same site. LCI-S enables identification of the initial sites of DAMP release, which subsequently reach maximal signal intensity for detection via TIRF microscopy. In the majority of pyroptotic cells (74 out of 100 cells analyzed), the release sites for IL-1β and Hmgb1 were spatially distinct rather than overlapping, with a mean physical separation of approximately 20 μm. Conversely, a minor fraction of the cells (*n* = 26/100) showed overlapping release peaks for IL-1β and Hmgb1 (Fig. 4c, d), indicating that Ninj1-dependent membrane rupture can occur adjacent to Gsdmd pores.

**Figure 4.**
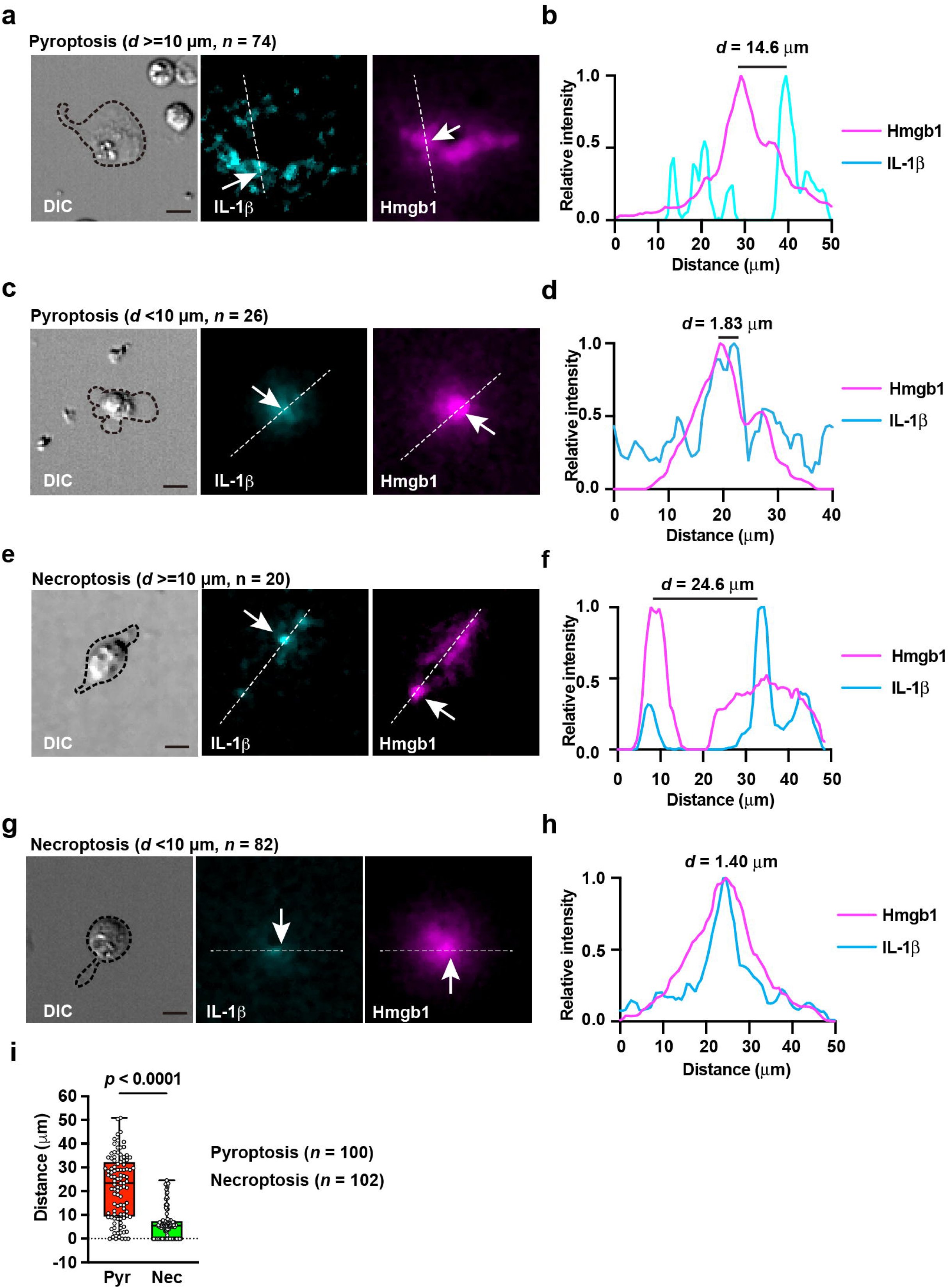
Distinct release sites of Hmgb1 and IL-1β during pyroptosis. Peritoneal macrophages were isolated, stimulated, and monitored by LCI-S as shown in Fig. 2. **a–h** Representative snapshot images showing the peak intensities of IL-1β and Hmgb1–mCherry release along with SYTOX uptake. The peak fluorescence positions for IL-1β and Hmgb1–mCherry were determined, and dotted lines were drawn between the two peaks (**a**, *n* = 74 cells; **b**, *n* = 26 cells; **f**, *n* = 20 cells; **g**, *n* = 82 cells). The fluorescence intensities along the dotted lines and the calculated distances between the two peak positions are shown (**b, d, f, h**). The peak intensity was normalized to 1.0. **i** Quantification of the distances between the peak release points of Hmgb1–mCherry and IL-1β during pyroptosis (*n* = 100 cells) and necroptosis (*n* = 102 cells). Box-and-whisker plots showing the median (centerline), interquartile range (box), and range (whisker), with each dot representing an individual cell. Pooled results from two independent experiments are shown. Statistical significance was assessed by the Mann–Whitney *U* test (**i**). Source data are provided as Supplementary Data 1.

In the vast majority of necroptotic cells (82 out of 102 cells analyzed), the peak fluorescence intensities of IL-1β and Hmgb1 exhibited nearly complete colocalization (Fig. 4e–i). Specifically, the distance between the most intense HMGB1–mCherry and IL-1β puncta was less than 5 µm. These findings are consistent with the fact that both HMGB1–mCherry and pro-IL-1β are released from membranes disrupted by MLKL activation.

### Spatiotemporal signaling profiles of distinct Hmgb1 release modes

It is inherently difficult to directly evaluate the biological consequences of distinct Hmgb1 release kinetics using conventional in vitro or in vivo experimental systems. To overcome this limitation, we applied a mathematical diffusion model based on Fick’s second law to quantitatively predict the spatiotemporal distribution of extracellular Hmgb1 ^13^. Because biological molecule release occurs over finite durations, we used the sustained release solution of the diffusion equation to model both short- and long-duration release ^13^. To establish biologically relevant model parameters, we first quantified the amount of Hmgb1 released from individual cells undergoing either pyroptosis or necroptosis via ELISA. Both cell types released approximately 4.3 ng mL^−1^ Hmgb1, corresponding to 11 fg cell^−1^ (Fig. 5a). In addition, we determined the minimum concentration of HMGB1 required to activate macrophages to be in the range of 30 to 100 ng mL^−1^ on the basis of inflammatory gene expression levels measured by quantitative PCR (Fig. 5b).

**Figure 5.**
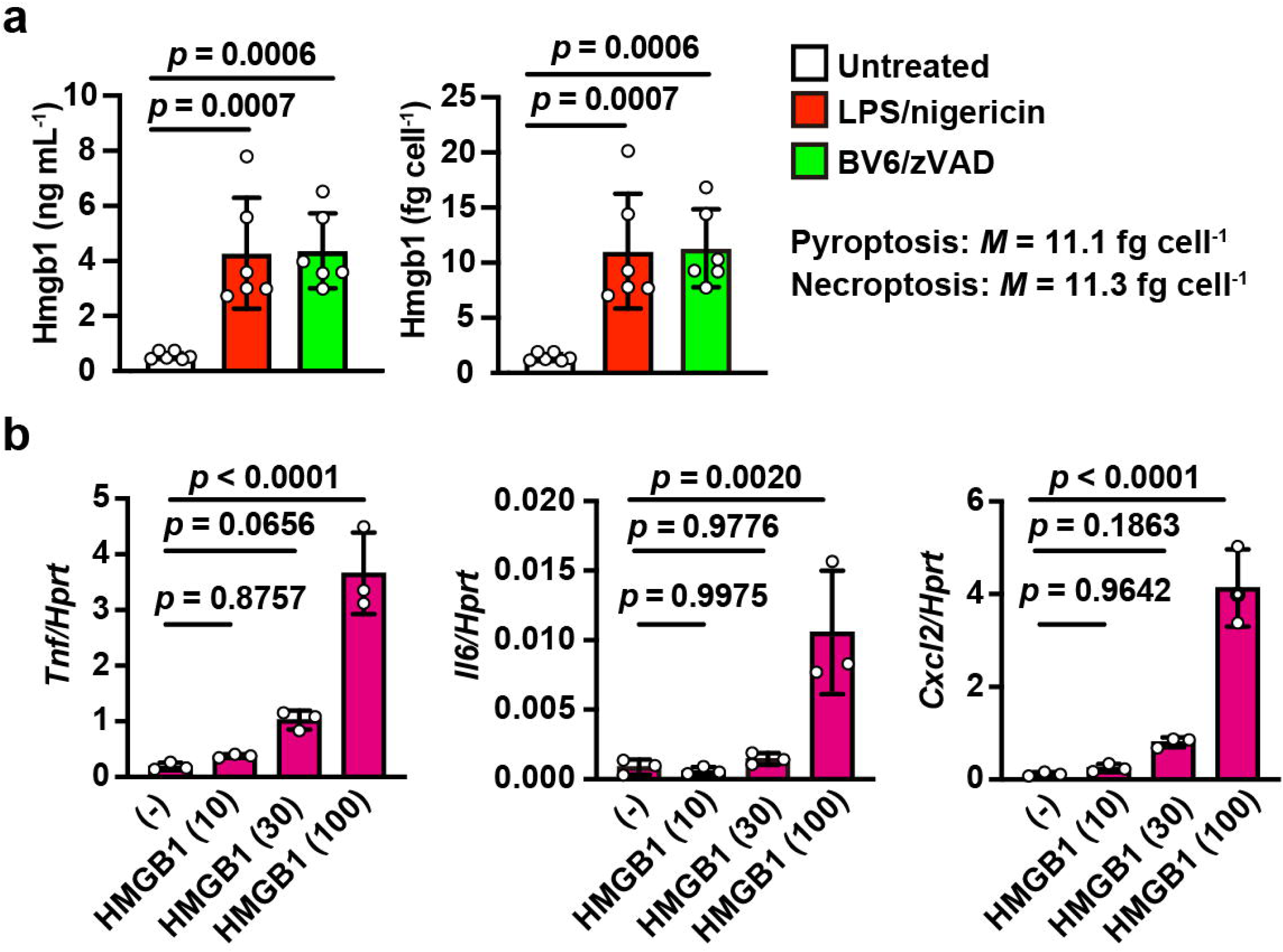
Spatiotemporal signaling profiles of distinct Hmgb1 release modes. **a** Peritoneal macrophages were prepared and stimulated as described in Fig. 1c, and HMGB1 concentrations in the culture supernatants were measured by ELISA. The absolute amount of Hmgb1 released per cell was calculated from these data using the equation described in the Methods section. The results are presented as mean ± SEM (*n* = 6). Each dot represents the value of cells from an individual mouse. The results are pooled from two independent experiments. **b** HMGB1 stimulates the expression of inflammatory cytokines and chemokines. Bone marrow-derived macrophages were prepared from wild-type mice and stimulated with the indicated concentrations (from 0 to 100 ng mL^−1^) of recombinant human HMGB1 for 4 h. The expression of the indicated genes was analyzed by qPCR. Data are presented as mean ± SD from three independent experiments, each performed in triplicate. Statistical analysis was performed using the one-way ANOVA with Dunnett’s multiple comparison test **(a, b**). Source data are provided as Supplementary Data 1.

The diffusion coefficient (*D*) of Hmgb1 was estimated to be approximately 100 µm² s^−1^ in water on the basis of its molecular weight and experimental measurements of proteins of comparable size ^20, 21^. Since extracellular matrix components and the viscosity of interstitial fluid reduce effective diffusion in vivo ^22, 23^, we used *D* = 40 µm² s^−1^ in the simulations to approximate physiological conditions (see the Methods section for details). We then calculated the spatiotemporal concentration profiles and gradients of Hmgb1 in each single cell at three representative distances. These distances were 20 µm, corresponding to local cell-to-cell signaling at approximately one to two cell diameters, and 50 and 100 µm, which are commonly used in in vitro chemotactic assays ^24, 25^. Compared to the long-duration mode, short-duration release at a 20 µm radius generated a 9-fold higher peak concentration, exceeding 30 ng mL□^1^, and a 12-fold steeper gradient, while reaching these maxima more than 10-fold earlier (Table 1). These results suggest that short-duration release efficiently induces strong local chemotactic responses and activates nearby macrophages. In contrast, at longer distances, the difference between the short-duration and long-duration release values was substantially smaller than that at 20 µm (Table 1). Similar results were obtained from the analyses of the release kinetics of Hmgb1 from necroptotic cells (Table 1).

**Table 1.** Distinct spatio-temporal signaling profiles of Hmgb1 release modes.

**Pyroptosis:** short ( $T = 0.31$ min) and long ( $T = 3.89$ min)
| Parameter | $r = 20 \mu\text{m}$ | | $r = 50 \mu\text{m}$ | | $r = 100 \mu\text{m}$ | |
| --- | --- | --- | --- | --- | --- | --- |
|  | Short | Long | Short | Long | Short | Long |
| Duration (min) | 0.31 | 3.89 | 0.31 | 3.89 | 0.31 | 3.89 |
| Time to Peak (min) | 0.32 | 3.89 | 0.40 | 3.93 | 0.87 | 4.12 |
| Max conc. ($\text{ng mL}^{-1}$) | 35.74 | 4.14 | 5.59 | 1.34 | 0.80 | 0.44 |
| Max grad ($\nabla C$) ($\text{ng mL}^{-1} \mu\text{m}^{-1}$) | 2.84 | 0.23 | 0.32 | 0.04 | 0.03 | 0.01 |
| Time at Max Grad (min) | 0.31 | 3.89 | 0.35 | 3.91 | 0.61 | 4.00 |
| Dur > 10% of max (min) | 0.82 | 5.34 | 2.29 | 8.11 | 7.98 | 13.88 |

**Necroptosis:** short ( $T = 0.30$ min) and long ( $T = 10.24$ min)
| Parameter | $r = 20 \mu\text{m}$ | | $r = 50 \mu\text{m}$ | | $r = 100 \mu\text{m}$ | |
| --- | --- | --- | --- | --- | --- | --- |
|  | Short | Long | Short | Long | Short | Long |
| Duration (min) | 0.30 | 10.24 | 0.30 | 10.24 | 0.30 | 10.24 |
| Time to Peak (min) | 0.31 | 10.24 | 0.39 | 10.27 | 0.87 | 10.41 |
| Max conc. ($\text{ng mL}^{-1}$) | 36.58 | 1.65 | 5.63 | 0.59 | 0.80 | 0.23 |
| Max grad ($\nabla C$) ($\text{ng mL}^{-1} \mu\text{m}^{-1}$) | 2.93 | 0.09 | 0.33 | 0.01 | 0.03 | 0.00 |
| Time at Max Grad (min) | 0.30 | 10.24 | 0.34 | 10.26 | 0.60 | 10.33 |
| Dur > 10% of max (min) | 0.80 | 12.30 | 2.28 | 16.62 | 7.97 | 24.44 |

Averages of each value at the indicated distances was derived from mathematical simulations based on the duration of Hmgb1 release from pyroptotic or necroptotic cells (see the Method section). The following parameters of Hmgb1 were used: $M = 11$ fg $\text{cell}^{-1}$ ; $D = 40 \mu\text{m}^2 \text{s}^{-1}$ .

### Recruitment of Gr-1^+^ Ly6C□ inflammatory monocytes in cisplatin-injured kidneys

Our in vitro and modeling results suggest that short-duration Hmgb1 release maximizes its local concentration to drive the activation of neighboring cells. Given that HMGB1 is known to facilitate immune cell recruitment via CXCL12 complex formation ^26, 27^, we next examined whether such local release events drive immune cell recruitment in vivo. Using two-photon intravital imaging of Hmgb1–mCherry transgenic mice, we visualized the dynamics of Hmgb1 release and immune cell behavior. Building on our previous intravital imaging model of necroptosis in a cisplatin-induced kidney injury model ^10^, we first identified the immune cell populations recruited to injured kidneys. Flow cytometric analysis revealed a marked increase in both the frequency and number of CD11b□ Gr-1□ cells but not CD11b□ F4/80□ cells following cisplatin administration (Fig. 6a–c, Supplementary Fig. 3a). This increase correlated with elevated serum levels of BUN and creatinine, as well as the upregulation of tissue injury markers (Fig. 6d, e). Immunohistochemistry confirmed the accumulation of Gr-1□ cells in the kidneys on Day 2 after cisplatin injection (Fig. 6f, g). These cells were also positive for the monocyte/macrophage marker CD68 (Supplementary Fig. 4). Phenotypic analysis revealed that most recruited Gr-1□ cells expressed Ly6C but not Ly6G (Fig. 6h–j, Supplementary Fig. 3b), indicating that these cells are inflammatory monocytes rather than neutrophils.

**Figure 6.**
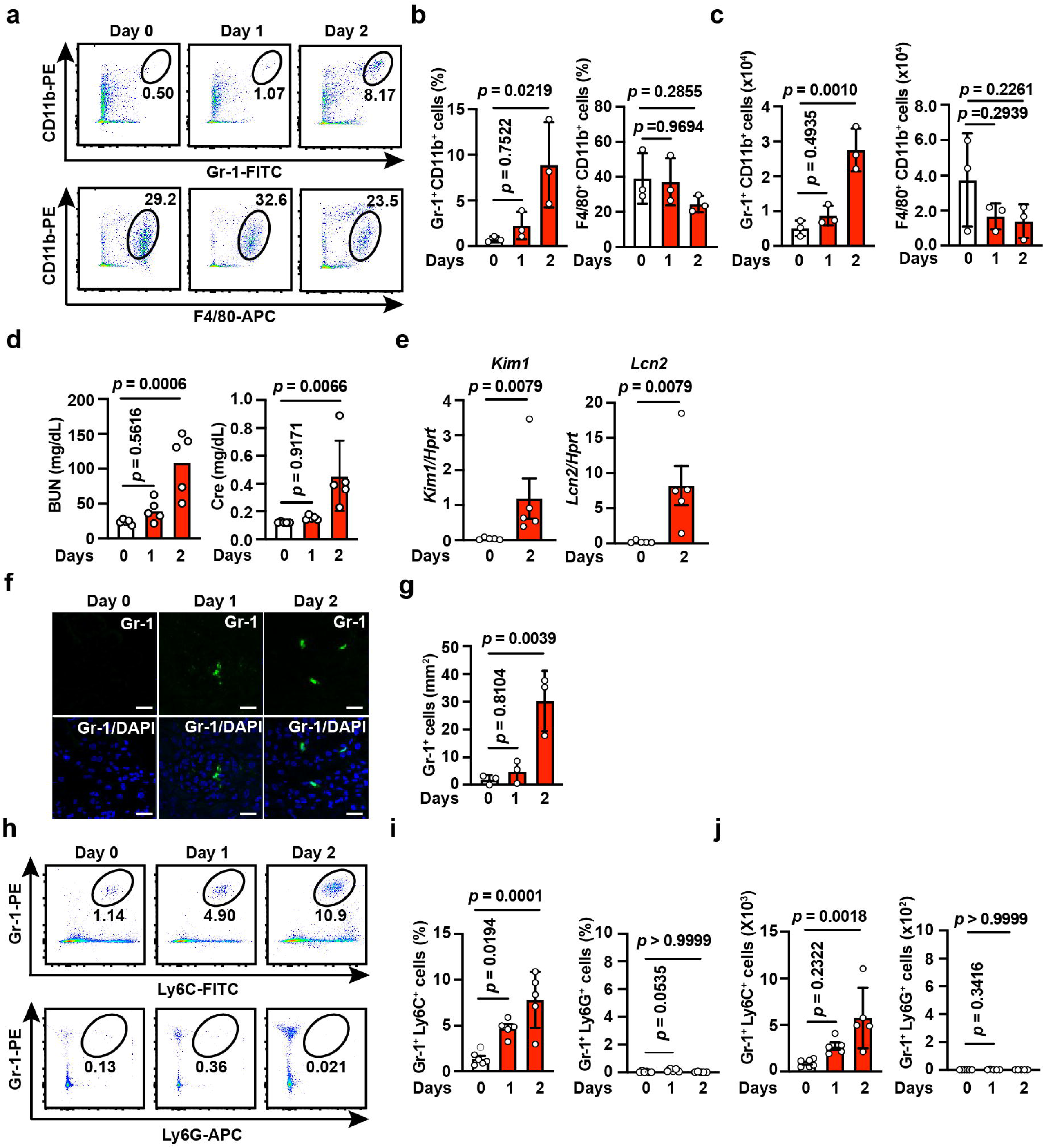
Cisplatin administration induces Gr-1□ Ly6C□ lymphocyte infiltration into the kidney. Eight-week-old wild-type mice were intraperitoneally injected with cisplatin (20 mg kg^−1^) and analyzed at the indicated time points. **a–c** Infiltration of Gr-1□ CD11b□ cells into the kidney following cisplatin treatment. Kidney mononuclear cells from untreated or cisplatin-treated mice were analyzed by flow cytometry. Representative FACS plots and the percentages of Gr-1□ CD11b□ and F4/80□ CD11b□ cells among CD45□ cells are shown (**a**). The percentages (**b**) and absolute numbers (**c**) of the indicated populations are summarized. Results are mean ± SEM (*n* = 3 mice/time point). **d** Increased serum BUN and Cre levels following cisplatin treatment. Serum BUN and Cre concentrations were measured. Data are presented as mean ± SEM (*n* = 5 mice/time point). **e** Increased expression of injury markers. The expressions of *Kim1* and *Lcn2* were quantified by qPCR. Data are presented as mean ± SEM (*n* = 5 mice/time point). **f, g** Gr-1□ cell accumulation in the kidney. Frozen kidney sections were stained with anti-Gr-1 antibody (**f**), and the Gr-1□ cells were quantified (**g**). Scale bar, 50 μm. Data are presented as mean ± SEM (*n* = 3 mice/time point). Representative images of three experiments are shown. **h–j** Infiltration of Gr-1□Ly6C□ cells after cisplatin administration. Kidney mononuclear cells were analyzed as described in (**a**). Representative FACS plots and the percentages of Gr-1□Ly6C□ and Gr-1□Ly6G□ cells among CD45□ cells are shown (**h**). The percentages (**i**) and absolute numbers (**j**) of each population are summarized. Statistical significance was determined by the one-way ANOVA with Dunnett’s multiple comparison test (**b, c, d, g, i-left, j**), Mann–Whitney *U* test (**e,** *Kim1*), Student’s *t*-test (**e**, *Lcn2*), or Kruskal-Wallis test with Dunnett’s multiple comparison test (**i-right**). Pooled results from three (**b, c, g**) or five (**d, e, i, j**) independent experiments are shown. Source data are provided as Supplementary Data 1.

### Gr-1□ cell infiltration following nuclear Hmgb1 depletion

HMGB1 expression in RTE cells markedly decreased after cisplatin treatment compared with that in the untreated controls, which is consistent with the release of Hmgb1 from damaged cells (Fig. 7a, b). To assess the dynamics of inflammatory monocyte recruitment in vivo, circulating Gr-1□ cells were fluorescently labeled by intravenous injection of Alexa Fluor 488-conjugated anti-Gr-1 antibody as described elsewhere ^28^ and imaged on Days 1 and 2 after cisplatin administration (Fig. 7c). Alexa Fluor 680-conjugated dextran was coinjected to visualize the capillary network, as cisplatin-induced injury disrupts vascular integrity and increases vascular permeability ^29^. In control mice, Hmgb1–mCherry fluorescence was uniformly distributed in RTE cell nuclei and the microvasculature interspersed among the tubules was intact, and indicated by the labeled dextran (Fig. 7d, left, Supplementary Movie 5). In contrast, dextran extravasated from the capillaries and blurred the capillary borders in cisplatin-treated mice as early as Day 1, an effect that became more pronounced by Day 2 (Fig. 7d, middle and right, Supplementary Movie 6). Quantitative analysis revealed progressive expansion of the dextran signal-positive area (Fig. 7e), which is indicative of structural remodeling associated with tissue damage.

**Figure 7.**
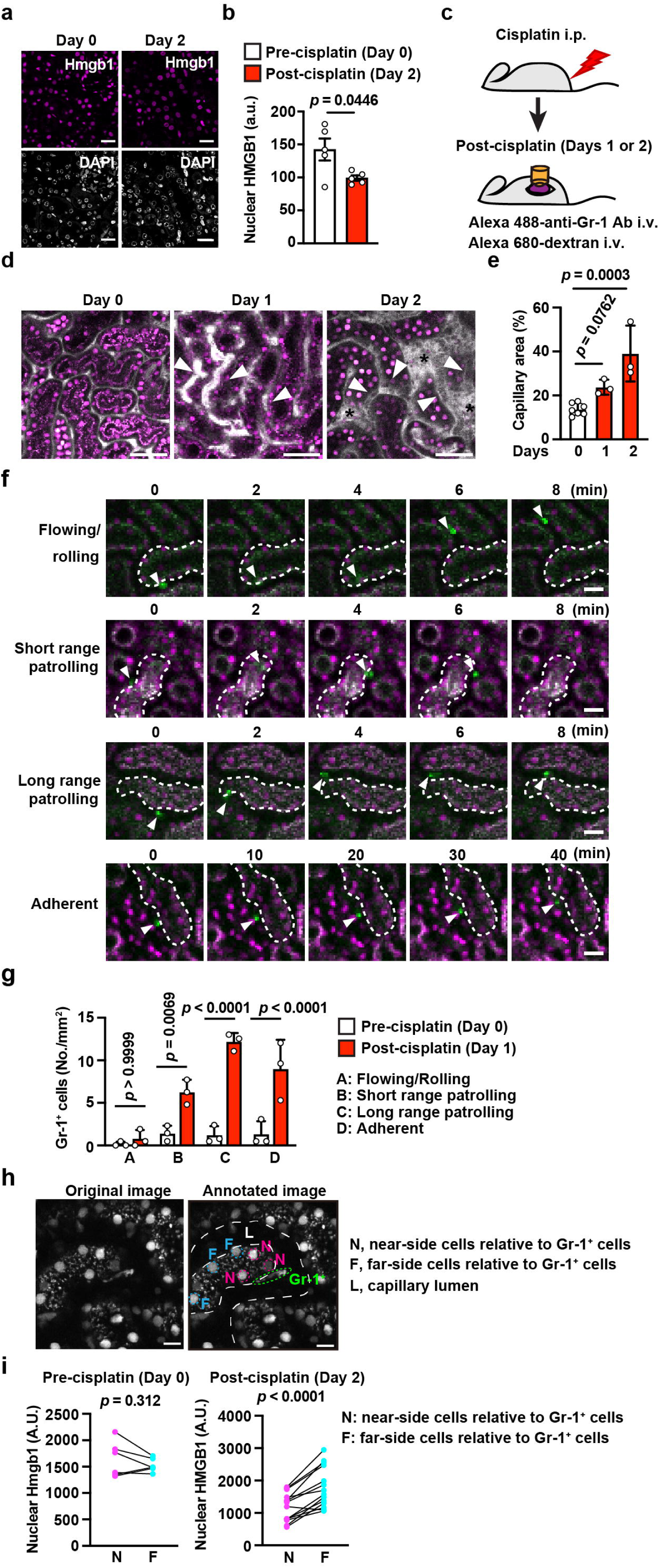
Gr-1□ cell infiltration following nuclear Hmgb1 depletion. **a, b** Decreased Hmgb1 fluorescence intensity in the kidney after cisplatin treatment. Eight-to twenty-week-old HMGB1–mCherry Tg mice were left untreated or injected intraperitoneally with cisplatin (20 mg kg^−1^). Frozen kidney sections, harvested at the indicated time points, were immunostained with anti-mCherry antibody. (**a**). Scale bar, 50 µm. Relative intensities of nuclear Hmgb1 were quantified and plotted (**b**). a.u.: arbitrary unit. Data are presented as mean ± SE (*n* = 5 mice/time point). **c–i** In vivo images of Hmgb1–mCherry Tg mice following cisplatin injection. **c** On Days 1 or 2 after treatment, the mice were anesthetized and positioned for two-photon intravital imaging. Alexa Fluor 488-conjugated anti-Gr-1 antibody and Alexa Fluor 680–dextran were administered immediately before imaging to visualize circulating Gr-1□ cells and capillary structures, respectively. Hmgb1–mCherry and Gr-1 signals were monitored over 3 h at 2-min intervals. **d** Representative images of HMGB1–mCherry fluorescence and Alexa Fluor 680–dextran distribution in the kidney are shown. White arrowheads indicate dilated capillaries; asterisks indicate extravasated dextran. Scale bar, 100 µm. **e**, The percentage of the capillary area in each field was quantified. Data are presented as mean ± SEM (Day 0, *n* = 7 mice; Day 1, *n* = 3 mice; Day 2, *n* = 3 mice) from three to four independent experiments. **f, g** Increased numbers of patrolling and adherent Gr-1□ cells in the kidney after cisplatin injection. Gr-1□ cell movements were tracked over time and categorized into four groups on the basis of the tracking rate and appearance frequency as previously described ^30^. Representative images of the four movement types are shown (**f**). White arrowheads indicate Gr-1□ cells (green), while magenta indicates Hmgb1–mCherry. Dotted lines outline the proximal tubules. Scale bar, 20 µm. The number of Gr-1□ cells exhibiting each movement type was quantified (**g**). Data are presented as mean ± SEM (*n* = 3 mice/time point). **h, i** Inverse relationship between nuclear Hmgb1 intensity in renal tubular epithelial (RTE) cells and the distance to Gr-1□ cells. Representative original image (left) and corresponding annotated image (right) showing Gr-1□ cells near Hmgb1–mCherry□ RTE cells at Day 2 after cisplatin injection (**h**). Scale bar, 20 µm. N, near-side cells; F, far-side cells; Gr-1^+^, Gr-1-positive monocytes. The white dotted lines indicate the capillary lumen (L). Nuclear Hmgb1 intensities were quantified in RTE cells located either near (N; ≤ 50 µm) or far from (F; > 50 µm) Gr-1□ cells in kidneys before cisplatin injection (Day 0, *n* = 6 cells) and 2 days after cisplatin injection (Day 2, *n* = 14 cells) (**i**). a.u., arbitrary unit. Pooled results from two independent experiments are shown. Statistical significance was determined by the two-tailed unpaired Student’s *t*-test (**b**), one-way ANOVA with Dunnett’s multiple comparison test (**e**), two-way ANOVA with Bonferroni’s multiple comparisons test (**g**), or Wilcoxon matched-pairs signed-rank test (**i**). Source data are provided as Supplementary Data 1.

Previous intravital imaging studies in the brain have classified monocyte motility into distinct states under ischemic conditions: flowing/rolling, short-range patrolling, long-range patrolling, and adherent ^30^. Similarly, few Gr-1□ cells were detected in the kidneys of the untreated mice because of their high circulation velocity (Fig. 7f, g, Supplementary Movie 7). In contrast, cisplatin exposure significantly increased the fractions of patrolling and adherent Gr-1□ monocytes, indicating increased retention and interaction within the microenvironment of the injured kidney (Fig. 7f, g). Of note, Gr-1^+^ monocytes engulfed RTE cells with diminished Hmgb1-mCherry signals and subsequently moved through the capillary lumen (Supplementary Movie 8). These observations suggest that RTE damage, accompanied by reduced Hmgb1–mCherry intensity, promotes the generation of chemoattractive cues—potentially including Hmgb1 itself—that mediate the recruitment of Gr-1□ monocytes and their subsequent engulfment of damaged cells.

To determine whether monocyte accumulation is spatially correlated with the extent of RTE damage, we quantified Hmgb1–mCherry intensity in RTE cells located both near and far from Gr-1□ cells (Fig. 7h). In untreated kidneys, Hmgb1–mCherry intensity was uniform regardless of the proximity to Gr-1□ cells (Fig. 7i, left). Conversely, in cisplatin-treated kidneys, compared with distant cells, RTE cells near Gr-1□ monocytes displayed significantly reduced Hmgb1–mCherry fluorescence (Fig. 7i, right). These findings establish an inverse spatial relationship between Hmgb1 signaling intensity and the localization of Gr-1□ monocytes, indicating that nuclear loss of Hmgb1 represents membrane rupture and the subsequent release of many DAMPs from cells as a cue for monocyte recruitment to injured RTE cells.

## Discussion

Using Hmgb1–mCherry transgenic mice, we visualized Hmgb1 release in vitro and in vivo. Simultaneous imaging with IL-1β revealed that Hmgb1 release was bimodal, whereas IL-1β release followed a single mode. Mathematical simulations indicated that short-duration Hmgb1 release from one or a few cells generated sufficient local concentrations to activate neighboring macrophages. In vivo imaging revealed that inflammatory monocytes were preferentially recruited to RTE cells characterized by a reduction in intracellular Hmgb1–mCherry signals.

Simultaneous imaging of Hmgb1–mCherry and IL-1β release from macrophages derived from Hmgb1–mCherry transgenic mice revealed that the bimodal release kinetics were specific to Hmgb1 and were not observed for IL-1β. Although a direct quantitative comparison of the release durations of Hmgb1 and IL-1β is precluded by the differences in their respective detection systems ^9, 11^, the lack of two distinct release clusters for IL-1β suggests that the bimodal pattern reflects an intrinsic property of Hmgb1 release. The relatively prolonged release of IL-1β from pyroptotic cells is mechanistically consistent with its reported two-step process: proteolytic maturation of pro-IL-1β by caspases is followed by IL-1β secretion through GSDMD pores ^16, 17^. In contrast, pro-IL-1β release from necroptotic cells might be expected to be more rapid, as it is passively released through MLKL-mediated membrane rupture without the need for proteolytic processing ^1^. Nevertheless, the mean release duration of pro-IL-1β from necroptotic cells also approached 100 minutes, which is strikingly slow even after accounting for the additional delay in detection inherent to the IL-1β imaging system ^11^. These findings suggest that an active regulatory mechanism may lead to the slow release of IL-1β that is independent of its maturation. Mathematical modeling to correct for differences in detection systems is necessary to precisely define and compare the true release kinetics of Hmgb1 and IL-1β. Supporting the notion that bimodal release kinetics may represent a unique feature of HMGB1, our recent study showed that IL-33 release exhibited a single short-duration mode in multiple IL-33–mCherry-expressing cell types ^31^.

In vivo labeling of Gr-1□ cells by intravenous injection of Alexa Fluor 488-conjugated anti-Gr-1 antibody enabled visualization of Gr-1□ cells circulating within the renal vasculature. Consistent with a previous study using a brain injury model ²□, the behavior of rolling Gr-1□ cells shifted toward short- or long-range patrolling patterns. Notably, we observed that some Gr-1□ rolling cells within capillary vessels abruptly extravasated and adhered to specific RTE cells in which the Hmgb1–mCherry intensity had declined. These observations suggest that damage signals released from injured epithelial cells can influence the behavior of circulating immune cells in vivo. In contrast to the in vitro injury model, it is likely in the in vivo injury model that multiple damaged cells contribute to the overall DAMP signal within the tissue. Nevertheless, the preferential recruitment of circulating Gr-1□ cells to particular injured epithelial cells suggests the presence of spatially localized signals that guide immune cell targeting. In this context, locally released DAMPs may directly contribute to immune cell recruitment and/or stimulate surrounding cells to produce chemokines that promote the further recruitment of Gr-1□ cells. Together, these findings raise the possibility that locally released DAMPs, potentially in combination with additional microenvironmental cues, contribute to the selective recruitment of immune cells to sites of tissue injury.

Our analyses revealed a trade-off between peak concentration and signal persistence during Hmgb1 release. Although long-duration release generates lower peak concentrations than short-duration release, it maintains concentrations greater than 10% of the peak at a distance of 20 μm for 6- to 15-fold longer. In vivo, where multiple cells often undergo death simultaneously, the reduced peak concentration associated with long-duration release may be offset by the cumulative release of Hmgb1 from multiple cells, enabling effective signal propagation within the local tissue environment. In addition, the prolonged persistence of Hmgb1 may favor its function as a chemoattractant. HMGB1 has been reported to form complexes with CXCL12 and promote immune cell recruitment ^7, 8^, suggesting that sustained rather than transient exposure may be critical for this activity. Consistent with this idea, the temporal advantage of long-duration release was evident at short distances (e.g., 20 µm) but decreased at longer distances (e.g., 50 or 100 µm), where diffusion dominates signal attenuation. These findings suggest that DAMPs primarily act within the immediate vicinity of dying cells, regardless of release kinetics. However, short- and long-duration release may result in distinct functional outputs: short-duration release may preferentially drive the rapid activation of nearby immune cells through high local concentrations and steep gradients, whereas long-duration release may support sustained chemotactic signaling through prolonged exposure. Together, our results raise the possibility that the kinetics of Hmgb1 release serve as a regulatory mechanism that shapes local immune responses during tissue injury. Further studies, including the development of approaches to quantify DAMP dynamics in vivo, will be needed to test this hypothesis.

Our present study provides a better understanding of the fundamental difference in how DAMPs and chemokines establish effective extracellular gradients. In contrast to the findings discussed above, chemokines such as CXCL8 participate in long-range leukocyte recruitment through a fundamentally different mechanism: they are continuously secreted by activated stromal and immune cells and are immobilized on extracellular glycosaminoglycans (GAGs), generating stable haptotactic gradients that extend over distances of 50–100 µm or more in tissue ^25, 32^. This ECM-bound gradient is essential for the directional guidance of neutrophils and monocytes from the vasculature into the tissue parenchyma, a process that operates on a spatial scale that far exceeds the effective signaling range of Hmgb1 released from a single cell.

A methodological caveat arises from the difference between Hmgb1 and IL-1β detection systems. Released Hmgb1**–**mCherry is directly captured by anti-mCherry-coated surfaces and visualized by TIRF microscopy, whereas released IL-1β requires sequential capture and detection with fluorescently labeled antibody. Consequently, the interval between the onset of SYTOX fluorescence and IL-1β signal detection is intrinsically longer than that observed for Hmgb1–mCherry, thereby precluding direct comparison of their release kinetics.

## Methods

### Reagents

We purchased the following reagents: cisplatin (AG-CR1-3590, AdipoGen), BV6 (B4653, APExBIO), FVD506 (65-0866-14, Invitrogen), GSK’872 (530389, Merck), human HMGB1 (C-terminal His Tag, 10326-H08H, Sino Biological), LPS (434, List Labs), nigericin (AG-CN2-0020, AdipoGen), murine TNF (34-8321, eBioscience), and zVAD (3188-v, Peptide Institute). The following antibodies were used in this study: anti-HMGB1 (ab18256, Abcam, diluted 1:1000), anti-mCherry (26765-1-AP, Proteintech, 1:5000), anti-GAPDH (2118S, Cell Signaling Technology, 1:1000), anti-GSDMD (ab209845, Abcam, 1:1000), anti-IL-1β (AF-401-NA, R&D, 1:1000), anti-MLKL (3H1, Millipore, 1:1000), anti-RIPK3 (63-216, ProSci, 1:1000), anti-phospho-MLKL (Ser345) (37333, Cell Signaling Technology, 1:1000), anti-phospho-RIPK3 (57220, Cell Signaling Technology, 1:1000), horseradish peroxidase (HRP)-conjugated donkey anti-rat IgG (712-035-153, Jackson ImmunoResearch Laboratories, Inc., 1:20000), HRP-conjugated donkey anti-rabbit IgG (NA934, GE Healthcare, 1:20000), biotin-conjugated donkey anti-rabbit IgG (E0432, Dako, 1:200), streptavidin-HRP (P0397, Dako, 1:300), APC-Cy7-anti-CD45 (157617, BioLegend, 1:500), phycoerythrin (PE)-anti-CD11b (50-0112-U100, TOMBO, 1:500), PE cyanine-7-anti-CD11b (101215, BioLegend, 1:500), FITC-anti-CD11b (101206, BioLegend, 1:500), APC-anti-F4/80 (20-4801-U100, TOMBO, 1:500), FITC-anti-F4/80 (123107, BioLegend, 1:500), PE-anti-Gr-1 (50-5931-U100, TOMBO, 1:500), FITC-anti-Gr-1 (35-5931-U100, TOMBO, 1:500), and FITC-anti-Ly6C (128005,BioLegend, 1:500).

### Cells

To isolate PEC macrophages, 6- to 8-week-old mice of the indicated genotype were intraperitoneally injected with 2.5 ml of 3% thioglycollate (T9032, Sigma). On Day 4 after thioglycollate injection, anesthetized mice were intraperitoneally injected with ice-cold PBS; then, the peritoneal cells were harvested with PBS. This procedure was repeated twice. Harvested cells were placed in plates containing RPMI medium. After nonadherent cells were removed, the attached cells were used as PEC macrophages. Approximately 80 to 90% of these cells were stained with antibodies against CD11b and F4/80. The cells were cultured in 10% fetal bovine serum (FBS)-containing RPMI and used for stimulation within one day. Cells were primed with LPS (10 ng mL^−1^) for 3 h and then stimulated with 5 μM or 20 μM nigericin for Western blotting and LCI-S, respectively, to induce pyroptosis. To induce necroptosis, cells were stimulated with BV6 (1 μM) and zVAD-fmk (20 μM) in the absence or presence of the RIPK3 inhibitor GSK’872 (5 μM).

Bone marrow cells were isolated from the femurs of 8-week-old C57BL/6 mice. Briefly, the bones were harvested, cleaned, and flushed with sterile PBS to extract the bone marrow. The obtained cell suspension was filtered through a 70-μm strainer, centrifuged at 300 × g for 5 minutes, and treated with RBC lysis buffer. After being washed, the cells were resuspended in DMEM supplemented with 10% FBS and 10 ng mL^-1^ recombinant mouse M-CSF and seeded at 1–2 × 10□ cells mL^-1^ in Petri dishes. The cells were cultured at 37°C with 5% CO□, and the medium was changed every 2–3 days. After 7 days, the adherent cells were harvested and used as bone marrow-derived macrophages (BMDMs) for subsequent experiments. BMDMs were stimulated with the indicated concentrations of recombinant human HMGB1 for 4 h and then subjected qPCR.

### Mice

Hmgb1-mCherry Tg mice were generated by microinjecting *Tol2* mRNA and the pT2KXIG-m Hmgb1–mCherry vector ^9^ into the cytoplasm of fertilized eggs from C57BL/6 mice ^33^. C57BL/6 mice were purchased from Sankyo Lab Service Co., Ltd.

The mice were housed in a specific pathogen-free facility under a 12-h light/dark cycle and provided free access to a routine chow diet and water. All animal experiments were performed according to the guidelines approved by Institutional Animal Care and Use Committees of Faculty of Medicine, Toho University (approval number: 23-530), Kyoto University Graduate School of Medicine (approval number: Medkyo 21562), and the National Cerebral and Cardiovascular Center (approval number: 25065).

### Western blotting

Murine tissues were homogenized with Polytron (Kinematica, Inc.) and lysed in RIPA buffer (50 mM Tris-HCl (pH 8.0), 150 mM NaCl, 1% Nonidet P-40, 0.5% deoxycholate, 0.1% SDS, 25 mM β-glycerophosphate, 1 mM sodium orthovanadate, 1 mM sodium fluoride, 1 mM phenylmethylsulfonyl fluoride, 1 μg/ml aprotinin, 1 μg/ml leupeptin, and 1 μg/ml pepstatin). Cells were lysed in RIPA buffer to generate whole-cell lysates. After centrifugation, tissue homogenates or cell lysates were subjected to SDS–polyacrylamide gel electrophoresis under reducing conditions, unless otherwise indicated, and transferred to polyvinylidene difluoride membranes (IPVH 00010, Millipore). In certain experiments, the same samples were subjected to SDS□PAGE under nonreducing conditions. The membranes were immunoblotted with the indicated antibodies and developed with Super Signal West Dura Extended Duration Substrate (34076, Thermo Scientific). The signals were analyzed with an Amersham Imager 600 (GE Healthcare Life Sciences).

### Flow cytometry

Cells were stained with the indicated antibodies in flow cytometry staining buffer (eBioscience). The prepared cells were gated on forward and side scatter to identify the lymphocyte population, followed by doublet discrimination. The cells were analyzed with a BD LSRFortessa X-20 flow cytometer (BD Biosciences) and FlowJo software (BD Biosciences).

### Live-cell imaging of secretion activity (LCI-S)

PEC macrophages isolated from Hmgb1–mCherry Tg mice were plated on a PDMS-glass hybrid microwell array chip precoated with anti-mCherry and anti-IL-1β antibodies. After the cells were primed with LPS (10 ng mL^−1^) for 3 h, the culture medium was replaced with fresh medium containing 1% bovine serum albumin (BSA). Subsequently, the cells were stimulated with nigericin (20 μM) to induce pyroptosis or with BV6 (1 μM) + zVAD (20 μM) to induce necroptosis. Mineral oil was layered on top of the medium to prevent evaporation. We observed the microwells at 2-minute intervals to detect intranuclear and intracellular Hmgb1–mCherry signals by epifluorescence microscopy and the extracellular release of Hmgb1–mCherry or IL-1β by TIRF-M. Visualization of Hmgb1–mCherry release by LCI-S was performed as previously described with some modifications ^34^. Briefly, time-resolved measurements were taken with a completely automated inverted microscope (ECLIPSE Ti-E; Nikon) equipped with a high numerical aperture (NA) objective lens (CFI Apo TIRF 60× Oil, NA=1.49; Nikon), a stage-top incubator (INUBG2TF-WSKM; Tokai Hit Co.) and an EM-CCD camera (ImagEM C9100-17; Hamamatsu Photonics K.K.). A high-pressure mercury lamp (Intensilight, Nikon) and an LED (540–600 nm, X-Cite XLED1; Excelitas Technologies Corp.) were used as the light sources. The following sets of excitation (Ex) and emission (Em) filters and a dichroic mirror (DM) were used. For HMGB1–mCherry, Ex: FF01-559/34-25, Em: FF01-630/69-25, and DM: FF585-Di01-25×36; these optical filters were purchased from Semrock. For IL-1β-CF660, Ex: ZET405/470/555/640x (Chroma Technology), Em: HQ-series barrier filter for Cy5 (Nikon), and DM: ZT561dcrb-UF2 (Chroma Technology). For SYTOX blue, High Signal-to-Noise BL Series (large FOV imaging with a Ti2 microscope) filter cube for CFP (Nikon) was used. The time course of each fluorescence signal was analyzed using NIS Elements 4.6 (Nikon), and the relative intensities were calculated by ImageJ software.

The steepness (*α*) of Hmgb1–mCherry release was estimated by fitting the following modified logistic function to the data:

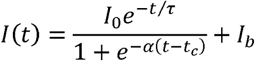

where *I*□ and *I_b_* are the maximum and basal intensities of the Hmgb1–mCherry release signal, respectively; τ is the time constant of exponential decrease due to mCherry photobleaching and Hmgb1–mCherry dissociation from the capture antibody; *α* is the steepness of the sigmoid curve; and *t*_c_ is the midpoint of the sigmoid. The duration *T* required for the signal to increase from 0.05 to 0.95 of the sigmoid curve was calculated as follows:

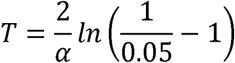

The experimentally derived values for *T* were subsequently used as input parameters for the spatiotemporal diffusion simulations. Only cells whose release profiles were well described by the modified logistic function (i.e., yielded a satisfactory goodness-of-fit) were included in further analyses.

For Hmgb1–mCherry or IL-1β release point analysis, the peak of the fluorescence signal in the captured image was detected, and the distance between the peak points was analyzed by ImageJ after time-lapse imaging.

### *k*-means clustering

Release durations were log-transformed and analyzed using *k*-means clustering ^19^. The number of clusters (*k*) was determined by integrating the statistical features of the distribution with biological interpretability. For Hmgb1, clustering with *k* = 2 led to clear separation in the log-transformed distribution, supporting classification into short- and long-duration release modes. Further subdivision into additional clusters was not pursued because the resulting groups lacked clear biological distinction. In contrast, the duration of IL-1β release did not clearly separate in terms of distribution and was therefore treated as a continuous variable without clustering.

### Measurement of the distance between Hmgb1–mCherry and IL-1β peaks

Immediately after cells became SYTOX-positive, LCI-S images were analyzed using ImageJ (NIH). Representative cells undergoing pyroptosis or necroptosis were selected based on SYTOX uptake and morphological features in DIC images. Local fluorescence intensity peaks corresponding to Hmgb1–mCherry and IL-1β signals were identified using the “Find Maxima” function with identical threshold parameters applied across all images. The detected peak positions were manually verified and are indicated by arrowheads in the images. To quantify the spatial relationship between the two DAMP release sites, a straight line (dotted line) was drawn connecting the peak positions of Hmgb1 and IL-1β. Fluorescence intensity along this line was measured using the “Plot Profile” function. Intensity profiles for each channel were extracted along the same line, and values were normalized to the maximum intensity within each profile (set to 1.0) for comparison. The positions of peak fluorescence were determined from the intensity profiles, and the distance (*d*) between Hmgb1 and IL-1β peaks was calculated as the linear distance between these two maxima along the drawn line. Distances were converted from pixels to micrometers based on the microscope calibration. Cells were categorized based on the distance between peaks (e.g., *d* ≥ 10 μm or *d* < 10 μm), and representative images are shown with arrowheads indicating peak positions and dotted lines indicating the measurement axis. Quantification was performed across multipl cells (pyroptosis, *n* = 100; necroptosis, *n* = 102), and the distribution of distances is presented in the graph.

### Quantitative polymerase chain reaction (qPCR)

Total RNA was purified from BMDMs or mouse kidneys using Sepasol-RNA I SuperG (09379-55; Nacalai Tesque). Complementary DNA (cDNA) was synthesized using the ReverTra Ace qPCR RT Kit (FSQ-101, Toyobo). Quantitative PCR was conducted on a QuantStudio 3 Real-Time PCR System (Thermo Fisher Scientific) using SYBR Green chemistry. Gene expression levels were normalized to those of the endogenous control *Hprt* and analyzed using QuantStudio Design & Analysis Software v2.6. The sequences of the primers used are listed in Supplementary Table 1.

### Enzyme-linked immunosorbent assay (ELISA)

PEC macrophages from 8-week-old C57BL/6 mice were seeded in 96-well plates at 5.8 × 10□ cells per well in 0.15 mL of medium. The cells were primed with LPS (10 ng mL^−1^) for 3 h and then stimulated with nigericin (5 μM) for 8 h. To induce necroptosis, the cells were stimulated with BV6 (1 μM)/zVAD (20 μM) for 8 h. Under these experimental conditions, more than 90% of the cells died upon stimulation for 8 h. The concentrations of Hmgb1 in the culture supernatants were determined by ELISA according to the manufacturer’s instructions (KE00343, Proteintech). Briefly, the culture supernatants were incubated with a precoated ELISA plate for 90 min at room temperature (RT). After the plate was washed three times with wash buffer, it was incubated with biotinylated detection antibody for 60 min at RT. After three additional washes with the wash buffer, the plate was incubated with the HRP conjugate, followed by development with TMB substrate. After determining the Hmgb1 concentration in the culture supernatants, the absolute amount of Hmgb1 per cell was calculated using the following equation:

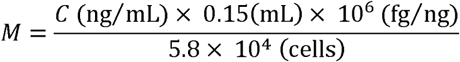

### Spatiotemporal diffusion simulations

To estimate the spatiotemporal distribution of Hmgb1 release during cell death, we modeled extracellular diffusion using an analytical solution to the three-dimensional diffusion equation with a sustained point source. Given that the molecular weight of Hmgb1 (25 kDa) is similar to that of GFP (27 kDa), the diffusion coefficient in water (*D*) was estimated to be approximately 100 μm² s^−1^ on the basis of experimental measurements of proteins of comparable size ^20, 21^. Since extracellular matrix components and the viscosity of interstitial fluid reduce effective diffusion in vivo ^22^ ^23^, a value of *D* = 40 μm² s^−1^ was adopted for all the simulations.

Hmgb1 release was approximated as a continuous, uniform source with a total amount of *M* = 11 fg per cell that was released over a finite duration *T* (min). Under this assumption, the release rate is *q* = *M/T*. For a point source releasing a total mass of *M* at a constant rate of *q* = *M/T* over release duration *T*, the extracellular concentration C(*r, t*) at radial distance *r* (μm) from the source and time *t* (min) is expressed as:

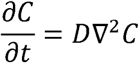

where *D* is the diffusion coefficient of Hmgb1 in the extracellular space and *t* and *T* are converted into seconds for all physical calculations.

The analytical solution for sustained release was obtained by temporal integration, yielding distinct expressions during and after the active release period.

During active release (*t* ≤ *T*):

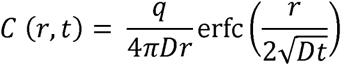

After cessation of release (*t* > *T*):

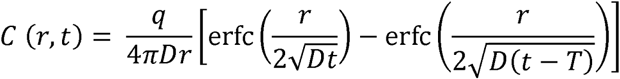

where erfc denotes the complementary error function defined as follows:

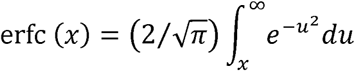

The concentrations were converted from fg μm^−3^ to ng mL^−1^ using a factor of 10^6^.

The peak concentration at distance *r* was determined by numerical search over the full time grid, as the time of peak concentration *t_p_*_eak_ depends on both the release duration *T* and the distance *r*. At short distances (*r* = 20 μm), *t*_peak_ occurs near the end of the release period (*t*_peak_ ≈ *T*), whereas at longer distances (*r* = 50 and 100 μm), diffusion delay causes *t*_peak_ to slightly exceed *T* (e.g., *t*_peak_ ≈ 0.40 min and 0.87 min, respectively, at *T* = 0.31 min).

To assess the chemotactic signaling capacity of Hmgb1 release, we calculated the spatial concentration gradient magnitude 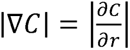 analytically.

Defining 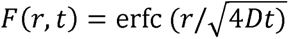 and its spatial derivative:

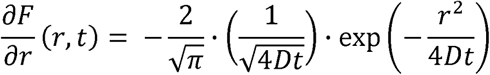

the gradient during active release (*t* ≤ *T*) is as follows:

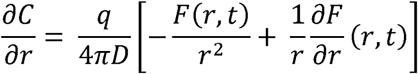

After cessation of release (*t* > *T*), the gradient becomes the following:

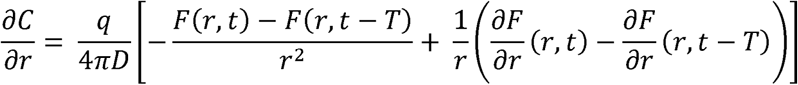

where *F*(*r,t − T*) and, ∂*F*/∂,*r*, (*r,t − T*) are evaluated at time (*t − T*).

All time-dependent quantities were evaluated over a logarithmically spaced time grid spanning 10^−6^ to 120 min (200,000 points). The peak concentration *C*_max_ and its time of occurrence *t*_peak_ were identified by numerical search over this grid at each distance *r*. The peak concentration was then recomputed precisely at *t*_peak_ using the analytical expression. The maximum spatial gradient |∇C|_max_ and its corresponding time were likewise determined by numerical search over the same time grid.

Analyses were performed at three representative distances: *r* = 20 μm, approximating the nearest-neighbor intercellular distance, and *r* = 50 and 100 μm, which are typically used in in vitro chemotactic assays ^24, 25^. Two cell death modalities were also examined: pyroptosis, with release durations of *T* = 0.31 min (short-duration) and *T* = 3.89 min (long-duration), and necroptosis, with *T* = 0.30 min (short-duration) and *T* = 10.24 min (long-duration). These release durations represent the mean values of each group, identified on the basis of the bimodal distribution of the live-cell imaging data of HMGB1 release kinetics. All calculations were implemented in Python (NumPy and SciPy libraries).

### Cisplatin-induced kidney injury model

Male C57BL/6 or Hmgb1–mCherry Tg mice (8–12 weeks old) were intraperitoneally injected with cisplatin (20 mg kg□^1^). Serum and kidney tissues were harvested at the indicated time points. Serum creatinine and blood urea nitrogen (BUN) levels were quantified by Oriental Yeast Co., Ltd. Detailed histological procedures are provided in the following subsections.

### Histological, immunohistochemical, and immunofluorescence analyses

The tissues were fixed in 10% formalin and embedded in paraffin blocks. Paraffin-embedded kidney sections were subjected to hematoxylin and eosin and immunohistochemical staining. For immunofluorescence analyses, frozen kidney sections were fixed with 4% formaldehyde and then stained with anti-mCherry, anti-CD68 or anti-Gr-1 antibodies, followed by Alexa Fluor 488-conjugated anti-rabbit or anti-rat antibodies in the presence of DAPI (1.25 μg mL^−1^). Images were acquired with a confocal microscope (LSM 880, Zeiss) and processed and analyzed with ZEN software (Zeiss) and the image processing package Fiji (https://fiji.sc/). The number of CD68□ or Gr-1□ cells was manually determined in four randomly selected microscopic fields.

### Isolation of mononuclear cells from kidneys

Kidneys were harvested from mice before or at the indicated time points after cisplatin injection. After decapsulation, the renal cortex was isolated and finely minced with surgical scissors. Tissue fragments were enzymatically digested in 1 mL of dissociation medium (calcium-containing HBSS) supplemented with collagenase (3 mg mL□^1^), DNase I (0.2 mg mL□^1^), and CaCl□ (50 µM), which had been prewarmed to 37 °C. The samples were gently stirred using a magnetic stirrer at 37 °C for 20 min, followed by mechanical dissociation using a 1-mL pipette tip; mechanical dissociation step was repeated twice at 5 min intervals. After digestion, the samples were diluted with 9 mL of ice-cold PBS containing 2% FBS, and cells were harvested by centrifugation at 1,500 rpm for 5 min. Mononuclear cells were then enriched by density gradient centrifugation using a 40%/80% Percoll gradient (800 × g, 20 min). Cells were recovered from the interphase and subsequently filtered through a 70-µm cell strainer.

### Quantification of Hmgb1-mCherry intensities of the kidney

DAPI-stained images were binarized using ImageJ to generate nuclear masks, which were used to define regions of interest (ROIs). mCherry IHC images were subjected to background subtraction in ImageJ, and the mean fluorescence intensity within each ROI was measured. The integrated signal intensity for each ROI was calculated as the product of the mean fluorescence intensity and the ROI area (mean × area), and the average value across the entire field of view was determined. The mean value from three independent fields was then calculated and used as a single data point per mouse. Quantification was performed using five mice per group, with or without cisplatin treatment, and the results are presented in the graph.

### Intravital imaging of mouse kidneys

Cisplatin (20 mg kg^−1^) was intraperitoneally injected into Hmgb1–mCherry Tg mice on Day 0. On Days 1 or 2, the mice were anesthetized using inhalational anesthesia, secured to a custom-made fixation platform, and placed under an objective lens for imaging. Alexa Fluor 488-conjugated anti-Gr-1 antibody (BioLegend, 108417, 3.5 µg per mouse) and Alexa Fluor 650–dextran (10,000 MW, anionic; Thermo Fisher, D34680, 100 µg per mouse) were intravenously injected to visualize circulating Gr-1□ cells and capillary lumens, respectively.

In vivo two-photon excitation microscopy was performed using an FVMPE-RS-SS-SP upright microscope (Evident) equipped with a 16×/0.6 objective (HC FLUOTAR L 16x/0.6 IMM CORR VISIR; Leica), an InSight X3 tunable ultrafast laser (SpectraPhysics), and an IR-cut filter (NDM750, Evident). The excitation wavelength for AF650 and mCherry was 1250 nm, a dichroic mirror (DM650, Evident) was used, as was an emission filter (BA660-750, Evident). AF650 emission is in the range of 660–750 nm, and that of mCherry is 575–645 nm; for Alexa Fluor 488, excitation is at 930 nm, a dichroic mirror (SDM475, Evident) was used, as was an emission filter (BA495-540, Evident). Images were acquired at different scan speeds depending on the depth. Once the images were combined into a single stack, the contrast was adjusted using the normalized layer function in Imaris software (Oxford). The acquired images were saved in multilayer 12-bit tagged image file format and processed and analyzed with MetaMorph software, as described previously ^35^.

### Statistic and Reproducibility

Statistical analyses were performed using GraphPad Prism (version 11.0.1; GraphPad Software). Data are presented as mean ± SD or SEM unless otherwise indicated. Statistical significance was determined using the unpaired two-tailed Student’s *t*-test, Mann□Whitney *U* test, one-way ANOVA with Dunnett’s or Tukey’s multiple comparisons test, Kruskal-Wallis test with Dunnett’s multiple comparison test, Wilcoxon matched-pairs signed-rank, or two-way ANOVA with Bonferroni’s multiple comparison test, as indicated in the figure legends. All experiments were independently repeated at least two or three times with similar results. Biological replicates represent independent experiments, whereas technical replicates refer to repeated measurements within the same experiment. Sample sizes (*n*), the number of independent experiments, and the statistical tests used for each dataset are provided in the corresponding figure legends. No statistical methods were used to predetermine sample size. The experiments were not randomized, and investigators were not blinded to allocation and outcome assessment.

## Acknowledgments

We thank K. Kawakami for providing the transposon-based expression vector system and acknowledge the technical support and imaging facilities provided by Kyoto University Live Imaging Center.

## Author Contributions

S.M., K.M., A.M., Ka.T. and Ke.T. and H.N. conceived and designed the study; S.M., D.-Y.S., A.M., Ka.T., M.Y. and S.K.-S. performed the experiments; K.S., M.Y., Y.S. and M.N. contributed new reagents and analytical tools; S.M., K.M., A.M., Ka.T., M.Y., Y.S., T.K., Ke.T. and H.N. analyzed the data; T.M., M.M. and K.O. supervised the study; and S.M. and H.N. wrote the manuscript with input from all the authors.

## Biological material availability

All the biological materials, including the Hmgb1–mCherry Tg mice used in this study, are available from the corresponding authors upon reasonable request.

## Data availability

All data supporting the findings of the study are available within the paper and its Supplementary Information. Source data underlying the main and Supplementary Figures are provided as Supplementary Data 1.

## Code availability

The Python code used for calculating the time to maximum concentration, maximum concentration, maximum gradient intensity, and time to maximum gradient intensity at distances of 20 μm, 50 μm, and 100 μm is provided in the Supplementary Code.

## AI usage

AI-assisted tools were used to support the development of the mathematical framework and Python code described in this study.

## References

1. Nakano H, Murai S, Moriwaki K. Regulation of the release of damage-associated molecular patterns from necroptotic cells. Biochem J 479, 677–685 (2022).

2. Ma M, Jiang W, Zhou R. DAMPs and DAMP-sensing receptors in inflammation and diseases. Immunity 57, 752–771 (2024).

3. Newton K, Strasser A, Kayagaki N, Dixit VM. Cell death. Cell 187, 235–256 (2024).

4. Jiang X, Stockwell BR, Conrad M. Ferroptosis: mechanisms, biology and role in disease. Nat Rev Mol Cell Biol 22, 266–282 (2021).

5. Tang D, Kang R, Zeh HJ, Lotze MT. The multifunctional protein HMGB1: 50 years of discovery. Nat Rev Immunol 23, 824–841 (2023).

6. Yang H, et al. MD-2 is required for disulfide HMGB1-dependent TLR4 signaling. J Exp Med 212, 5–14 (2015).

7. Yun J, et al. The HMGB1-CXCL12 Complex Promotes Inflammatory Cell Infiltration in Uveitogenic T Cell-Induced Chronic Experimental Autoimmune Uveitis. Front Immunol 8, 142 (2017).

8. Cecchinato V, et al. Redox-Mediated Mechanisms Fuel Monocyte Responses to CXCL12/HMGB1 in Active Rheumatoid Arthritis. Front Immunol 9, 2118 (2018).

9. Murai S, et al. A FRET biosensor for necroptosis uncovers two different modes of the release of DAMPs. Nat Commun 9, 4457 (2018).

10. Murai S, et al. Generation of transgenic mice expressing a FRET biosensor, SMART, that responds to necroptosis. Commun Biol 5, 1331 (2022).

11. Shirasaki Y, et al. Real-time single-cell imaging of protein secretion. Sci Rep 4, 4736 (2014).

12. Yamagishi M, Shirasaki Y. Live-Cell Imaging Technique to Visualize DAMPs Release During Regulated Cell Death. Methods Mol Biol 2274, 337–352 (2021).

13. Crank J. The mathematics of diffusion. Oxford University Press (1975).

14. Holditch SJ, Brown CN, Lombardi AM, Nguyen KN, Edelstein CL. Recent Advances in Models, Mechanisms, Biomarkers, and Interventions in Cisplatin-Induced Acute Kidney Injury. Int J Mol Sci 20, (2019).

15. Landau SI, et al. Regulated necrosis and failed repair in cisplatin-induced chronic kidney disease. Kidney Int 95, 797–814 (2019).

16. Kayagaki N, et al. Caspase-11 cleaves gasdermin D for non-canonical inflammasome signalling. Nature 526, 666–671 (2015).

17. Shi J, et al. Cleavage of GSDMD by inflammatory caspases determines pyroptotic cell death. Nature 526, 660–665 (2015).

18. Kayagaki N, et al. NINJ1 mediates plasma membrane rupture during lytic cell death. Nature 591, 131–136 (2021).

19. Wani AA. Comprehensive analysis of clustering algorithms: exploring limitations and innovative solutions. PeerJ Comput Sci 10, e2286 (2024).

20. Tyn MT, Gusek TW. Prediction of diffusion coefficients of proteins. Biotechnol Bioeng 35, 327–338 (1990).

21. Milo R, Phillips R. Cell biology by the numbers. Garland Science, Taylor & Francis Group (2016).

22. Pluen A, et al. Role of tumor-host interactions in interstitial diffusion of macromolecules: cranial vs. subcutaneous tumors. Proc Natl Acad Sci U S A 98, 4628–4633 (2001).

23. Sykova E, Nicholson C. Diffusion in brain extracellular space. Physiol Rev 88, 1277–1340 (2008).

24. Penzo M, et al. Inhibitor of NF-kappa B kinases alpha and beta are both essential for high mobility group box 1-mediated chemotaxis [corrected]. J Immunol 184, 4497–4509 (2010).

25. Sarris M, et al. Inflammatory chemokines direct and restrict leukocyte migration within live tissues as glycan-bound gradients. Curr Biol 22, 2375–2382 (2012).

26. Schiraldi M, et al. HMGB1 promotes recruitment of inflammatory cells to damaged tissues by forming a complex with CXCL12 and signaling via CXCR4. J Exp Med 209, 551–563 (2012).

27. Mantonico MV, et al. The acidic intrinsically disordered region of the inflammatory mediator HMGB1 mediates fuzzy interactions with CXCL12. Nat Commun 15, 1201 (2024).

28. Finsterbusch M, et al. Patrolling monocytes promote intravascular neutrophil activation and glomerular injury in the acutely inflamed glomerulus. Proc Natl Acad Sci U S A 113, E5172–5181 (2016).

29. Gu J, et al. The disruption and hyperpermeability of blood-labyrinth barrier mediates cisplatin-induced ototoxicity. Toxicol Lett 354, 56–64 (2022).

30. Masthoff M, et al. Resolving immune cells with patrolling behaviour by magnetic resonance time-lapse single cell tracking. EBioMedicine 73, 103670 (2021).

31. Kanokogi T, et al. Temporal control of Ninj1 activation determines cell-to-cell Heterogeneity in IL-33 release. Communications Biology in press, (2026).

32. Proudfoot AEI, Johnson Z, Bonvin P, Handel TM. Glycosaminoglycan Interactions with Chemokines Add Complexity to a Complex System. Pharmaceuticals (Basel) 10, (2017).

33. Sumiyama K, Kawakami K, Yagita K. A simple and highly efficient transgenesis method in mice with the Tol2 transposon system and cytoplasmic microinjection. Genomics 95, 306–311 (2010).

34. Liu T, et al. Single-cell imaging of caspase-1 dynamics reveals an all-or-none inflammasome signaling response. Cell Rep 8, 974–982 (2014).

35. Kamioka Y, et al. Live imaging of protein kinase activities in transgenic mice expressing FRET biosensors. Cell Struct Funct 37, 65–73 (2012).

