## Supplementary Materials for "Hmgb1 release kinetics shape its extracellular functions during regulated cell death"

**Supplementary Figures 1.** Gating strategy of peritoneal macrophages.

**Supplementary Figures 2.** Clustering of Hmgb1 and IL-1β release durations.

**Supplementary Figures 3.** Gating strategy of mononuclear cells prepared from kidneys of untreated or cisplatin-treated mice.

**Supplementary Figures 4.** Infiltration of $CD68^+$, but not $Ly6G^+$ cells, in the cisplatin-treated kidneys.

**Supplementary Figures 5.** Uncropped images of the Western blots shown in Fig. 1a, 1c.

**Supplementary Table 1.** Primers used in this study.

**Supplementary Movie 1.** Short-duration Hmgb1 and IL-1β release during pyroptosis. Time 0 indicates the start of imaging. Scale bar, 20 μm.

**Supplementary Movie 2.** Long-duration Hmgb1 and IL-1β release during pyroptosis. Time 0 indicates the start of imaging. Scale bar, 20 μm.

**Supplementary Movie 3.** Short-duration Hmgb1 and IL-1β release during necroptosis. Time 0 indicates the start of imaging. Scale bar, 20 μm.

**Supplementary Movie 4.** Long-duration Hmgb1 and IL-1β release during necroptosis. Time 0 indicates the start of imaging. Scale bar, 20 μm.

**Supplementary Movie 5.** Mouse kidney on Day 0, Alexa Fluor 650–dextran injection. Time 0 indicates the start of imaging. Scale bar, 100 μm.

**Supplementary Movie 6.** Mouse kidney on Day 2 after cisplatin injection, Alexa Fluor 650–dextran injection. Time 0 indicates the start of imaging. Scale bar, 100 μm.

**Supplementary Movie 7.** Movement of Gr-1$^+$ cells in the mouse kidney on Day 0 visualized with Alexa Fluor 488-conjugated anti-Gr-1 antibody. Time 0 indicates the start of imaging. Scale bar, 20 μm.

**Supplementary Movie 8.** Movement of Gr-1$^+$ cells in the mouse kidney on Day 2 after cisplatin treatment visualized with Alexa Fluor 488-conjugated anti-Gr-1 antibody. Time 0 indicates the start of imaging. Scale bar, 20 μm.

**Supplementary code**

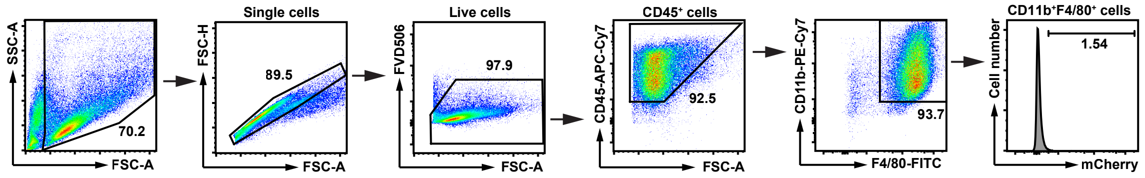


**Supplementary Figure 1 | Gating strategy of peritoneal macrophages, related to Fig. 1.** Peritoneal macrophages prepared from 8- to 17-week-old wild-type and Hmgb1-mCherry Tg mice were stained with FVD506, APC-Cy7-anti-CD45, PE-Cy7-anti-CD11b, and FITC-anti-F4/80 antibodies. The cells were analyzed with flow cytometry. Results are representative of five independent experiments. The percentage of the cell populations in each panel is shown.

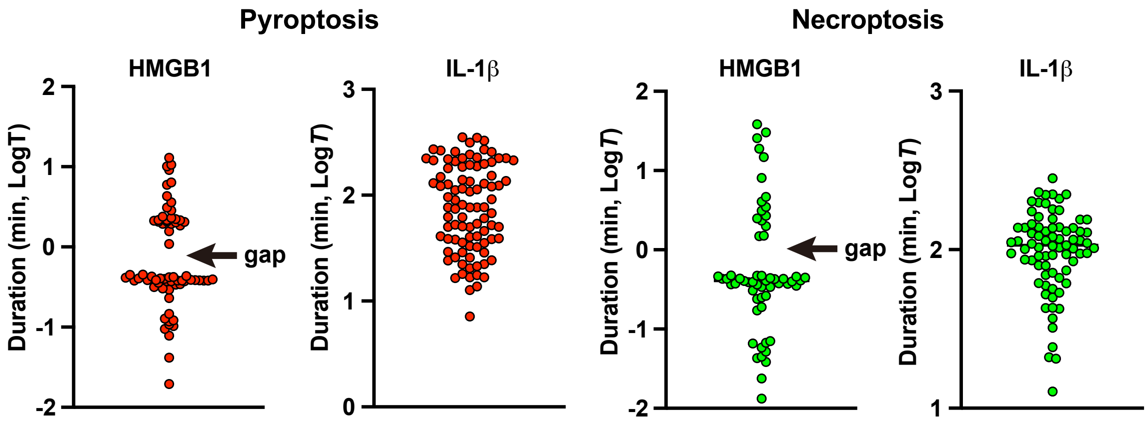


**Supplementary Figure 2 | Clustering of Hmgb1 and IL-1β release durations, related to Figs. 2 and 3**. The duration of Hmgb1 and IL-1β release from individual cells undergoing pyroptosis and necroptosis was calculated as described in the Methods and plotted on a logarithmic scale (y-axis). Notably, gaps were observed in Hmgb1 release, but not in IL-1β release, in both forms of cell death. Source data are provided as Supplementary Data 1.

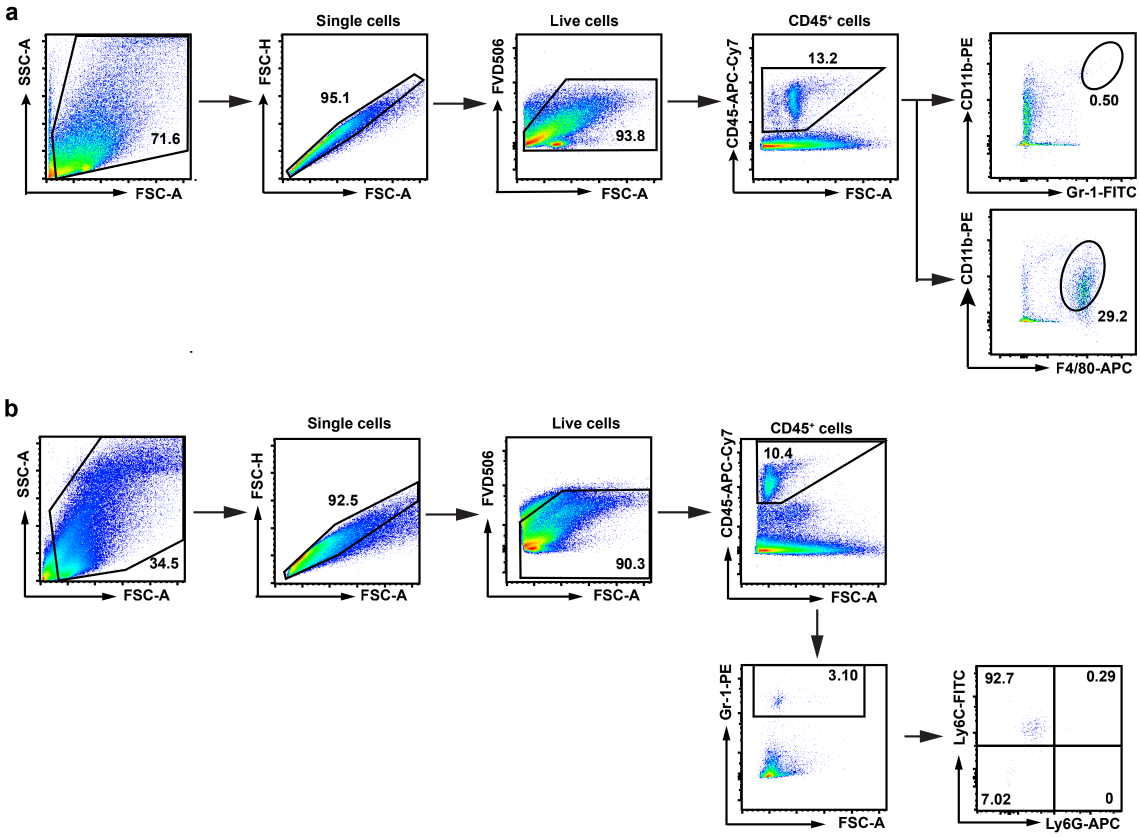


**Supplementary Figure 3 | Gating strategy of mononuclear cells prepared from kidneys of untreated or cisplatin-treated mice, related to Fig. 6. a, b** Eight-week-old wild-type mice were untreated or intraperitoneally injected with cisplatin as in Fig. 6. $CD45^+$ cells were prepared from the kidneys by collagenase perfusion, followed by Ficoll density gradient centrifugation as described in the Methods. Cells were stained with FVD506, APC-Cy7-anti-CD45, PE-anti-CD11b, and APC-anti-F4/80 (**a**) or FVD506, APC-Cy7-anti CD45, PE-anti-Gr-1, FITC-anti-Ly6C and APC-anti-Ly6G (**b**). The cells were analyzed with flow cytometry. Results are representative of four independent experiments. The percentage of the cell populations in each panel is shown.

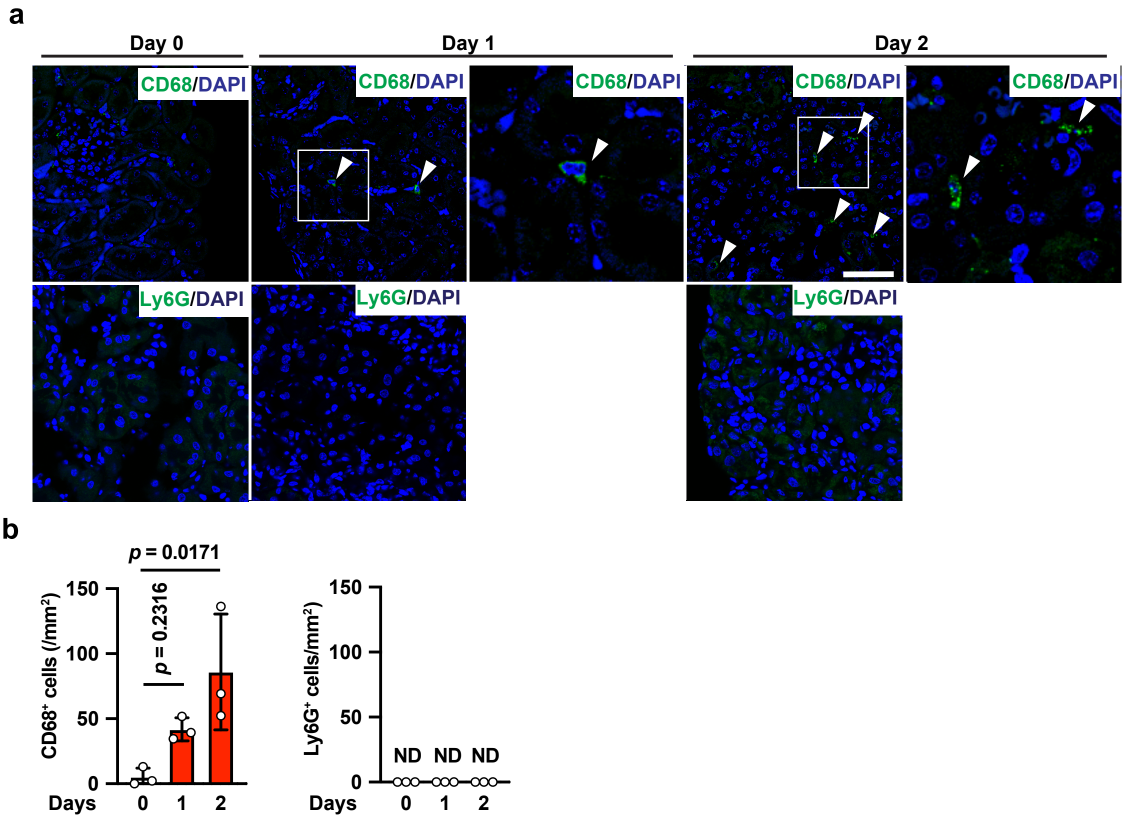


**Supplementary Figure 4 | Infiltration of CD68$^+$, but not Ly6G$^+$ cells, in the cisplatin-treated kidneys, related to Fig. 6.** Eight-week-old wild-type mice were intraperitoneally injected with cisplatin as in Fig. 5. **a** Frozen tissue sections were stained with anti-CD68 or anti-Ly6G antibodies, and the immunofluorescent signals were analyzed by confocal microscopy. Right panels on Day 1 and Day 2 are enlarged images of the left boxes. The white arrowheads indicate CD68$^+$ cells. Scale bar, 50 µm. **b** The number of CD68$^+$ or Ly6G$^+$ cells were counted by randomly picking up high power field ($n = 3$ mice). ND, not detected. Pooled Results of two independent experiments. Statistical significance was determined by the one-way ANOVA with Dunnett's multiple comparison test. Source data are provided as Supplementary Data 1.

**Supplementary Figure 5; uncropped images of the Western blots shown in Figs.1a, 1c**

**Fig.1a**

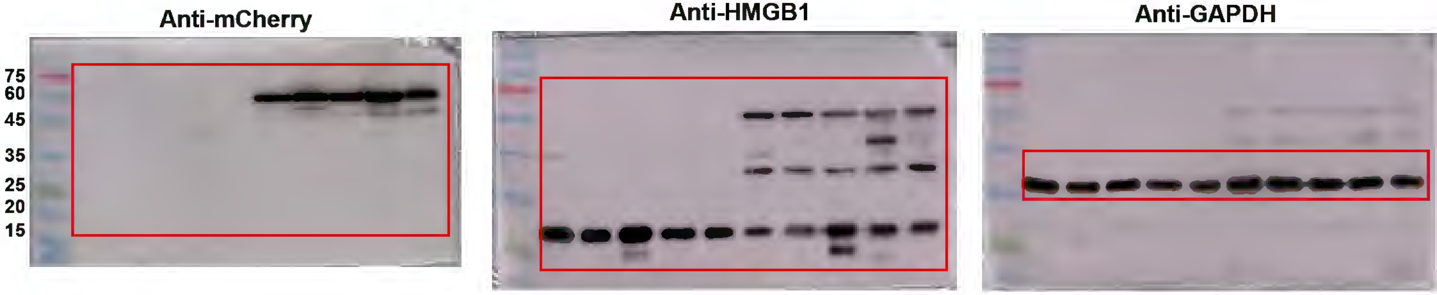


**Fig.1c**

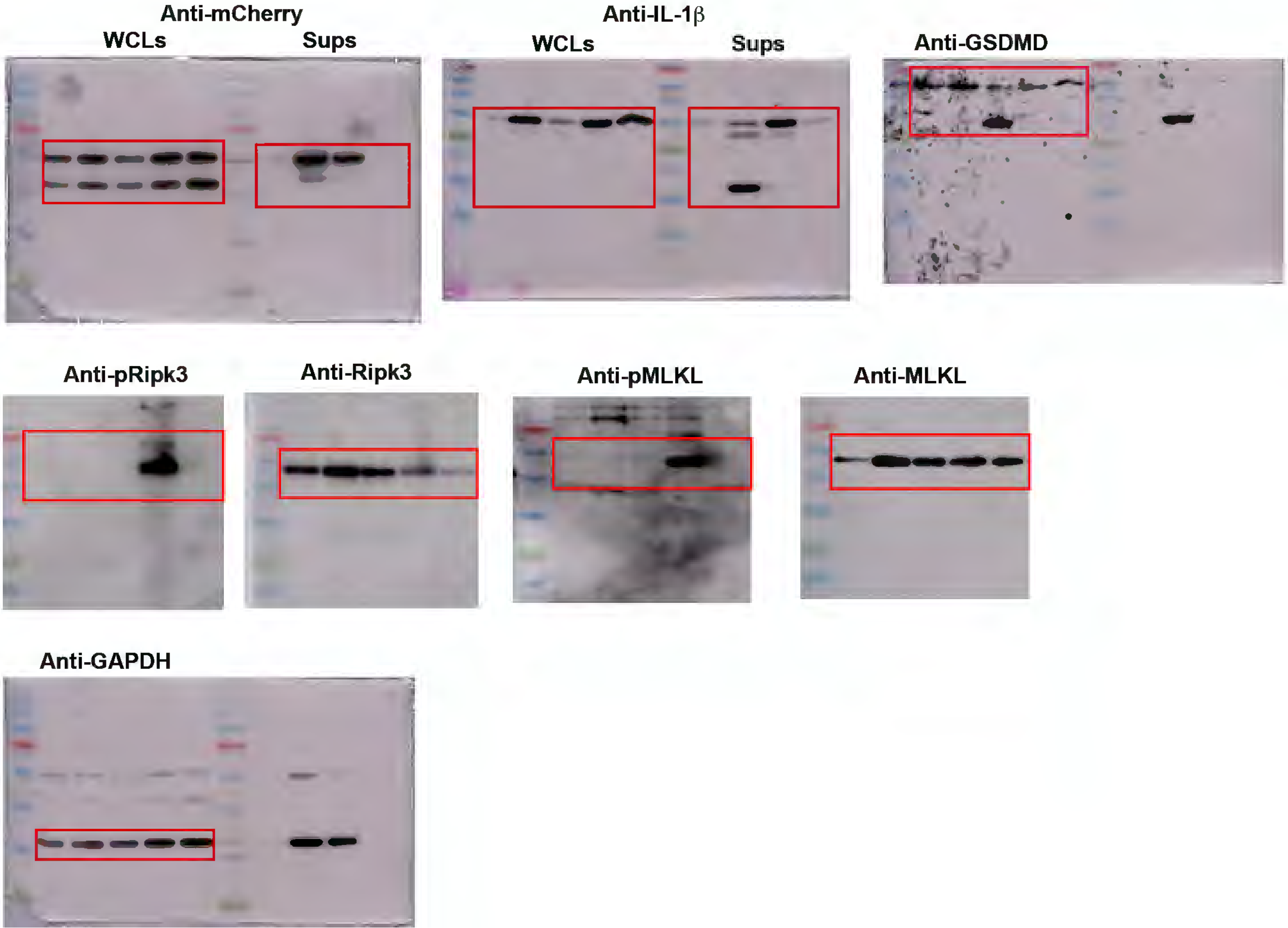

**Supplementary Table 1. Primers used in this study.**

| | | |
|---|---|---|
| *Cxcl2:* | Fwd | 5’- CCAACCACCAGGCTACAGG-3’ |
| | Rev | 5’- GCGTCACACTCAAGCTCTG-3’ |
| *Hprt:* | Fwd | 5’-AACAAAGTCTGGCCTGTATCCAA-3’ |
| | Rev | 5’-GCAGTACAGCCCCAAAATGG-3’ |
| *Il6*: | Fwd | 5’- GTATGAACAACGATGATGCACTTG-3’ |
| | Rev | 5’- ATGGTACTCCAGAAGACCAGAGGA-3’ |
| *Kim1* | Fwd | 5’- GTCTGTATTGTTGTCGAGTGGAG-3’ |
| | Rev | 5’- CGTGTGGGAATCTCTGGTTTAAC-3’ |
| *Lcn* | Fwd | 5’- AAGGAGCTGTCCCCTGAACT-3’ |
| | Rev | 5’- GGTGGGGACAGAGAAGATGA-3’ |
| *Tnf*: | Fwd | 5’- GAAAAGCAAGCAGCCAACCA-3’ |
| | Rev | 5’- CGGATCATGCTTTCTGTGCTC-3’ |

**Supplementary code**

```
import numpy as np
import pandas as pd
from scipy.special import erfc

# ========================
# Parameters
# ========================
M = 11.0      # fg/cell
D = 40.0      # um^2/s (in vivo tissue condition)
R_LIST = [20.0, 50.0, 100.0]   # um
UNIT = 1e6     # fg/um^3 -> ng/mL
EPS = 1e-12
T_GRID = np.logspace(np.log10(1e-6), np.log10(120.0), 200000)   # 
min

CELL_TYPES = {
    "Pyroptosis":  {"short": {"T": 0.31}, "long": {"T": 3.89}},
    "Necroptosis": {"short": {"T": 0.30}, "long": {"T": 10.24}},
}


# ============================================
# Sustained release concentration
# ============================================
def C_sust(r, tmin, Tmin):
    """Extracellular  Hmgb1  concentration  (ng/mL)  at  distance  r 
(um)
    and  time  t  (min)  for  sustained  release  over  duration  Tmin 
(min)."""
    ts = np.maximum(np.asarray(tmin, dtype=float) * 60.0, EPS)
    Ts = Tmin * 60.0
    q = M / Ts
    pref = q / (4 * np.pi * D * r)
    term1 = erfc(r / np.sqrt(4 * D * ts))
```

```
    C_fg = np.empty_like(ts)
    on = ts <= Ts
    C_fg[on] = pref * term1[on]
    dt = np.maximum(ts[~on] - Ts, EPS)
    term2 = erfc(r / np.sqrt(4 * D * dt))
    C_fg[~on] = pref * (term1[~on] - term2)
    return C_fg * UNIT


# ======================================
# Concentration gradient: |∇C| = |∂C/∂r|
# ======================================
def dCdr_sust(r, tmin, Tmin):
    """Spatial concentration gradient |∂C/∂r| (ng/mL per um)
    for sustained release."""
    ts = np.maximum(np.asarray(tmin, dtype=float) * 60.0, EPS)
    Ts = Tmin * 60.0
    A = (M / Ts) / (4 * np.pi * D)
    out_fg = np.empty_like(ts)

    def F_and_Fp(time_s):
        a = 1.0 / np.sqrt(4.0 * D * time_s)
        arg = r * a
        return erfc(arg), -(2.0 / np.sqrt(np.pi)) * a * np.exp(-
(arg ** 2))

    on = ts <= Ts
    if np.any(on):
        F1, F1p = F_and_Fp(ts[on])
        out_fg[on] = A * (-F1 / r ** 2 + F1p / r)
    if np.any(~on):
        tso = ts[~on]
        F1o, F1op = F_and_Fp(tso)
        dt = np.maximum(tso - Ts, EPS)
        F2, F2p = F_and_Fp(dt)
        out_fg[~on] = A * (-(F1o - F2) / r ** 2 + (F1op - F2p) /
```

```
r)

    return out_fg * UNIT


# ========================
# Peak time (numerical search)
# ========================
def t_peak_sust_min(r, Tmin):
    """Time (min) at which concentration peaks at distance r
(um)."""
    C = C_sust(r, T_GRID, Tmin)
    return float(T_GRID[np.argmax(C)])


# ========================================
# Duration above fraction of own peak (min)
# ========================================
def duration_above_fraction(tmin, y, frac):
    """Total duration (min) where y(t) >= frac * max(y).
    Crossing times are estimated by linear interpolation."""
    y = np.asarray(y, dtype=float)
    t = np.asarray(tmin, dtype=float)
    thr = frac * np.max(y)
    above = y >= thr
    if not np.any(above):
        return 0.0
    idx = np.where(np.diff(above.astype(int)) != 0)[0]
    crossings = []
    for i in idx:
        t0, t1, y0, y1 = t[i], t[i + 1], y[i], y[i + 1]
        if abs(y1 - y0) < 1e-20:
            tc = 0.5 * (t0 + t1)
        else:
            tc = t0 + (thr - y0) * (t1 - t0) / (y1 - y0)
        crossings.append(tc)
```

```
    if above[0]:
        crossings = [t[0]] + crossings
    if above[-1]:
        crossings = crossings + [t[-1]]
    intervals = [
        (crossings[k], crossings[k + 1])
        for k in range(0, len(crossings), 2)
    ]
    return float(sum(b - a for a, b in intervals))


# ============================================
# Core analysis for a single (r, Tmin) pair
# ============================================
def analyze(r, Tmin):
    """Return key diffusion parameters at distance r for release
    duration Tmin."""
    t = T_GRID
    C = C_sust(r, t, Tmin)
    tp = t_peak_sust_min(r, Tmin)
    Cmax = float(C_sust(r, tp, Tmin))
    gs = np.abs(dCdr_sust(r, t, Tmin))
    ig = int(np.argmax(gs))
    return {
        "Duration (min)":           Tmin,
        "Time to Peak (min)":       tp,
        "Max conc. (ng/mL)":        Cmax,
        "Max grad (∇C) (ng/mL/um)": float(gs[ig]),
        "Time at Max Grad (min)":   float(t[ig]),
        "Dur > 10% of max (min)":   duration_above_fraction(t, C,
0.10),
    }


# ============================================
# Run analysis for all cell types and distances,
```

```
# and print results in a table.
# ==========================================
if __name__ == "__main__":
    rows = []
    for cell_type, modes in CELL_TYPES.items():
        for mode_name, params in modes.items():
            Tmin = params["T"]
            for r in R_LIST:
                res = analyze(r, Tmin)
                rows.append({
                    "Cell type": cell_type,
                    "Mode":      mode_name,
                    "r (um)":    r,
                    **res,
                })

    df = pd.DataFrame(rows)
    pd.set_option("display.float_format", lambda x: f"{x:.4f}")
    pd.set_option("display.width", 200)
    pd.set_option("display.max_columns", None)
    print(df.to_string(index=False))
```
